# Decoding and Reprogramming Redox Partner Specificity in Rieske Oxygenases for Enhanced Catalytic Activity

**DOI:** 10.64898/2026.04.03.713453

**Authors:** Hui Miao, Rick Oerlemanns, Peter-Leon Hagedoorn, Sandy Schmidt

## Abstract

Multicomponent Rieske oxygenases catalyze diverse oxidative transformations but require precisely matched redox partners to sustain efficient electron transfer, severely limiting their modularity and biocatalytic application. Yet, the molecular logic underlying this specificity remains poorly defined. Here we decode the molecular principles governing redox partner specificity in representative three-component Rieske oxygenase systems. Through systematic mutagenesis analysis and cross-component reconstitution assays, we identify a single ferredoxin residue that acts as a class-defining determinant of oxygenase recognition. Guided by this insight, we reprogram electron transfer between non-cognate components by complementary engineering of the oxygenase interface, creating an unnatural redox chain with substantially enhanced catalytic turnover compared to the native system. Spectroscopic, binding and computational analyses reveal that productive electron transfer arises from optimized electrostatic complementarity and redox potential alignment rather than maximal binding affinity. Extending this strategy to another oxygenase system demonstrates its generality. Together, these results establish transferable design rules for rationally engineering electron transfer pathways in multicomponent oxygenases, enabling their predictable adaptation as customizable biocatalysts.

---

Rieske oxygenases (ROs) constitute a large and functionally diverse family of non-heme iron enzymes that catalyze key oxidative transformations in both catabolic and anabolic pathways across bacteria^1^, animals^2^, and plants^3^. More than 70,000 members of this enzyme class have been identified, highlighting their widespread biological importance and catalytic versatility^4,5^. ROs catalyze a wide spectrum of chemo- and stereoselective reactions on aromatic and aliphatic substrates^6^, including hydroxylation and *N*-oxygenation^7^, as well as oxidative desaturation^2^, dealkylation^8–10^, and C–C bond formation or cleavage^11,12^. Despite their diverse chemistry, all ROs share a conserved catalytic framework comprising a [2Fe–2S] Rieske cluster and a mononuclear non-heme iron center within the α subunit^13^. These two cofactors operate in concert to mediate multi-electron transfer events necessary for O₂ activation and substrate oxidation^13^.

Similar to cytochrome P450s, ROs rely on electron transfer (ET) from redox partners to drive catalysis (Figure 1A). Based on their ET chain, ROs are classified into three major classes: the two-component class I systems composed of a reductase (Red) and an oxygenase (Oxy); and the three-component class II and III systems, which use an additional ferredoxin (Fd) as an intermediary electron carrier. The primary distinction between class II and class III systems lies in the nature of their Red components. Class II ROs can be further divided into subclasses IIA and IIB according to the type of Fd involved. Within these modular architectures, the efficiency and fidelity of inter-protein ET are key determinants of catalytic turnover, product yield, and protein stability. For example, hybrid cumene dioxygenase (CDO) systems incorporating more efficient non-native Red have shown enhanced ET efficiency and improved catalytic performance^14^.

**Figure 1.**
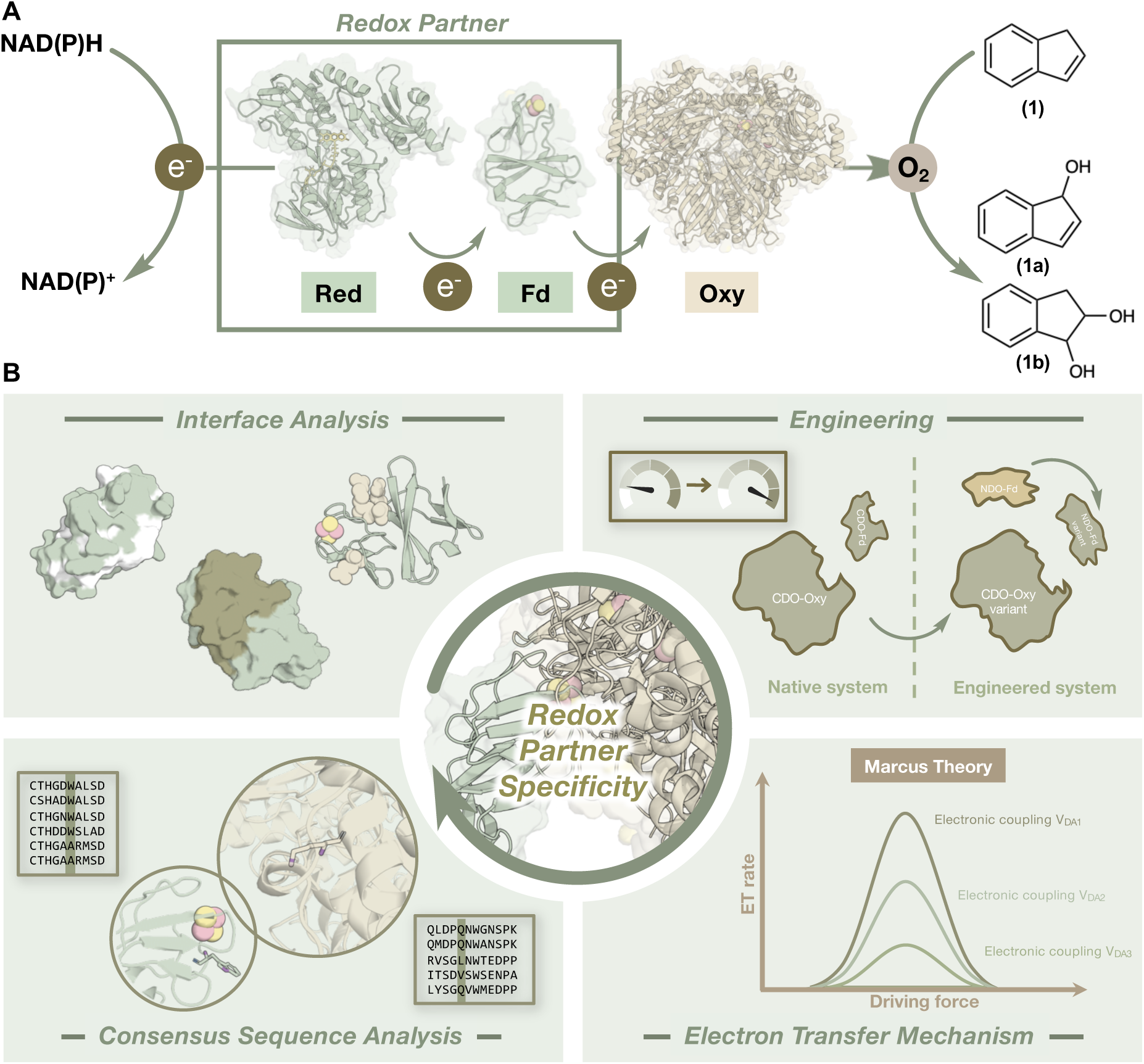
Electron transfer in three-component Rieske oxygenases (ROs) and reprogramming strategy employed in this study. (A) Overview of a typical three-component RO system, illustrated using cumene dioxygenase (CDO) from *P. fluorescens* IP01. The RO activity is driven by an electron transport chain (ETC) that utilizes NAD(P)H as the initial electron source. A reductase (Red) initiates the ETC by transferring electrons from NAD(P)H to a ferredoxin (Fd), which subsequently delivers single electrons to the terminal oxygenase component (Oxy). (B) Summary of the reprogramming strategy employed in this study, comprising interface analysis, rational protein engineering, analysis of electron transfer mechanism and consensus sequence analysis.

Despite the central role of ET for efficient catalysis, the molecular basis of redox partner recognition and specificity remains poorly understood in ROs. Even though the interchangeability of the Red in two-component RO systems has been reported^15–17^, clear differences in the activity of the Oxy with native and non-native Reds was observed^16,17^. Several studies performed with three-component ROs, such as the α_3_-type carbazole dioxygenase (CARDO), revealed that interactions between the terminal Oxy and its cognate Fd are typically highly specific, and substitution with non-native partners often lead to catalytic inefficiency^18^. Structural analysis of CARDO suggested that this specificity arises from finely tuned electrostatic, hydrophobic, and geometric complementarity at the protein–protein interface^19,20^. More recent docking and electrostatic potential analyses have extended these findings to α_3_β_3_-type Oxys, proposing conserved charge-based interactions with Rieske-type Fds^21^. Nevertheless, the principles that govern how redox partners in ROs can be tuned to be mutually compatible remain elusive. Such stringent specificity, while critical for efficient ET and thus catalysis, poses practical challenges for both mechanistic investigation and applied biocatalysis, as native redox partners are often unstable, difficult to express, or yet to be identified^4^. To address these challenges, attempts have been made to employ the hydrogen peroxide (H_2_O_2_) shunt pathway to bypass the need for the native redox partners^22^; however, this approach frequently yields low catalytic turnover and poor stability^23^. Furthermore, current efforts to identify compatible redox partners rely heavily on labor-intensive and time-consuming screening methods rather than rational design^17^. Therefore, elucidating the molecular determinants of redox partner compatibility is essential not only for deepening our mechanistic understanding of multi-component enzyme systems, but also for enabling the rational engineering of more catalytically efficient enzymes for biotechnological and industrial applications.

Redox partner engineering has achieved notable success in cytochrome P450 systems that are also dependent on ET mediated by redox partners, and offers a valuable framework for analogous strategies in ROs^24–29^. Optimization of protein–protein interactions and ET pathways in P450s through directed evolution, computational modeling, and mutagenesis has led to significant improvements in catalytic efficiency, including enhanced coupling and higher turnover rates^24,25^. For example, re-engineering the interface between the flavin mononucleotide (FMN) and heme domains of P450_BM3_ improved catalytic efficiency by more than 10-fold^26^. Furthermore, it was shown that the distance and orientation between redox centers are critical factors in ET efficiency^27,28^. For example, these insights have informed the redesign of PetF/OleP ET system to improve the biosynthesis of murideoxycholic acid^29^. Despite the clear similarities between P450s and ROs in their reliance on redox partner interactions, systematic interface engineering and ET pathway optimization have yet to be applied to the RO family.

In this study, we address this gap by investigating the molecular determinants that control redox partner specificity in representative RO systems (Figure 1B). We focus on CDO (class IIB) and naphthalene dioxygenase (NDO, class III) as model enzymes. Through mutagenesis analysis and cross-component reconstitution assays, we identified key residues—specifically, A50 in NDO-Fd and W48 in CDO-Fd—that define redox partner recognition and govern ET efficiency. Guided by these findings, we engineered the CDO-Oxy component further to establish an unnatural NDO–CDO ET pathway. Of the 17 designed variants, the CDO-Oxy mutant β_L98K paired with NDO-Red and NDO-Fd A50W exhibited a 3-fold increase in turnover number (TON_20min_) compared to the native CDO system. Mechanistic studies using EPR spectroscopy, binding affinity analysis, and docking simulations revealed that the reconstituted compatibility originated from a finely tuned redox potential gradient and optimized protein–protein interactions. Finally, we demonstrate that the identified redox partner recognition principles are conserved across other ROs, providing a general framework for modulating redox partner compatibility. These findings lay the groundwork for the rational design of versatile ROs with customized ET pathways, enabling their broader application as tunable redox biocatalysts.

## RESULTS

### Identification of Key Residues in Ferredoxin Determining Compatibility with Oxygenase

To elucidate the molecular basis of redox partner specificity in ROs, we selected the CDO and NDO systems as representative models. Both enzymes are three-component ROs^13^, comprising a Red, a Fd, and an Oxy, yet they exhibit considerable sequence divergence, only 41% identity between their Fd components, and 32% and 23% identity, respectively, for the α- and β-subunits of their respective Oxy. This evolutionary diversity provides an ideal framework to investigate how sequence variation translates into redox partner compatibility and differential ET efficiency. Each component was heterologously expressed in *Escherichia coli* JM109 (DE3) with an N-terminal His tag and purified according to established protocols^30^.

First, we evaluated the cross-reactivity of the two systems in *in vitro* assays using the conversion of indene **1** into 1*H*-indenol **1a** and 1,2-indandiol **1b** as the model reaction (Figure 1A, Scheme S1). CDO-Oxy showed negligible activity when paired with NDO-Fd, while NDO-Oxy retained partial activity with CDO-Fd (Figure S1), suggesting that CDO is more stringent in redox partner selection than NDO. Guided by the crystal structure of the Class III CARDO Fd–Oxy complex and a homology model of its Class IIB counterpart^20^, we selected candidate residues in CDO-Fd and NDO-Fd according to two criteria. First, we used sequence alignments to identify positions corresponding to Class III Fd residues that form electrostatic or hydrophobic contacts with the Oxy in the binary complex. These positions were E52, E62 and K87 in CDO-Fd and D42, T46, H47, D54, E63, L66, H67, P82 as well as K88 in NDO-Fd (Figure S2). Second, we included additional surface-exposed, unconserved charged and hydrophobic residues (E25, D41, R42, D47, W48 and E84 in CDO-Fd and R60, E61 as well as D72 in NDO-Fd) that could plausibly contribute to Fd–Oxy recognition specificity, even though they were not previously implicated. These residues were subjected to alanine-scanning mutagenesis to assess their functional relevance (Figure S3).

In CDO-Fd, alanine substitutions at positions D41, R42, E62, and K87 resulted in decreased product formation after 3 h, while the W48A mutation nearly abolished activity (Figure S3A). In contrast, the corresponding residues in NDO-Fd differ markedly in identity and polarity: D41 and R42 in CDO-Fd are replaced by asparagine and leucine, respectively. E63 and K88 correspond to E62 and K87 in CDO-Fd; however, mutagenesis revealed distinct effects. E63A unexpectedly enhanced activity, whereas K88A had minimal impact (Figure S3B). Among the residues investigated in NDO-Fd, mutations at H47 and H67, the ligands that coordinate the Rieske cluster, as well as L66, which corresponds to a cysteine in respiratory Rieske proteins that forms a disulfide bridge^31^, led to decreased product formation (Figure S3B). Due to the critical role of W48 in CDO-Fd, the reciprocal A50W mutation was introduced into NDO-Fd, which also nearly abolished activity (Figure S3B).

To further delineate how these residues shape Fd–Oxy interactions, we examined a series of targeted substitutions in CDO-Fd and NDO-Fd. In CDO-Fd (Figure 2A), the E62K variant partially restored activity relative to E62A, suggesting that E62 plays a role in hydrogen-bonding interaction (Figure 2B). A similar trend was observed for residue K87: the mutation to glutamic acid resulted in an almost inactive variant (K87E)), while variant K87Q retained moderate activity, indicating that this residue contributes to both electrostatic and hydrogen-bonding interactions (Figure 2B). In contrast, the D41R substitution completely disrupted activity, while the D41N substitution retained lower activity than the D41A substitution (Figure 2C). This indicates that D41 mediates critical electrostatic interactions with the Oxy surface. In addition, the indispensable role of W48 was underscored by the drastic loss of activity in the conservative W48F variant, which retained only ∼25% of wild-type activity (Figure 2D). Intriguingly, the E84Q and E84A mutation enhanced activity, likely due to the removal of an unfavorable electrostatic interaction (Figure 2E).

**Figure 2.**
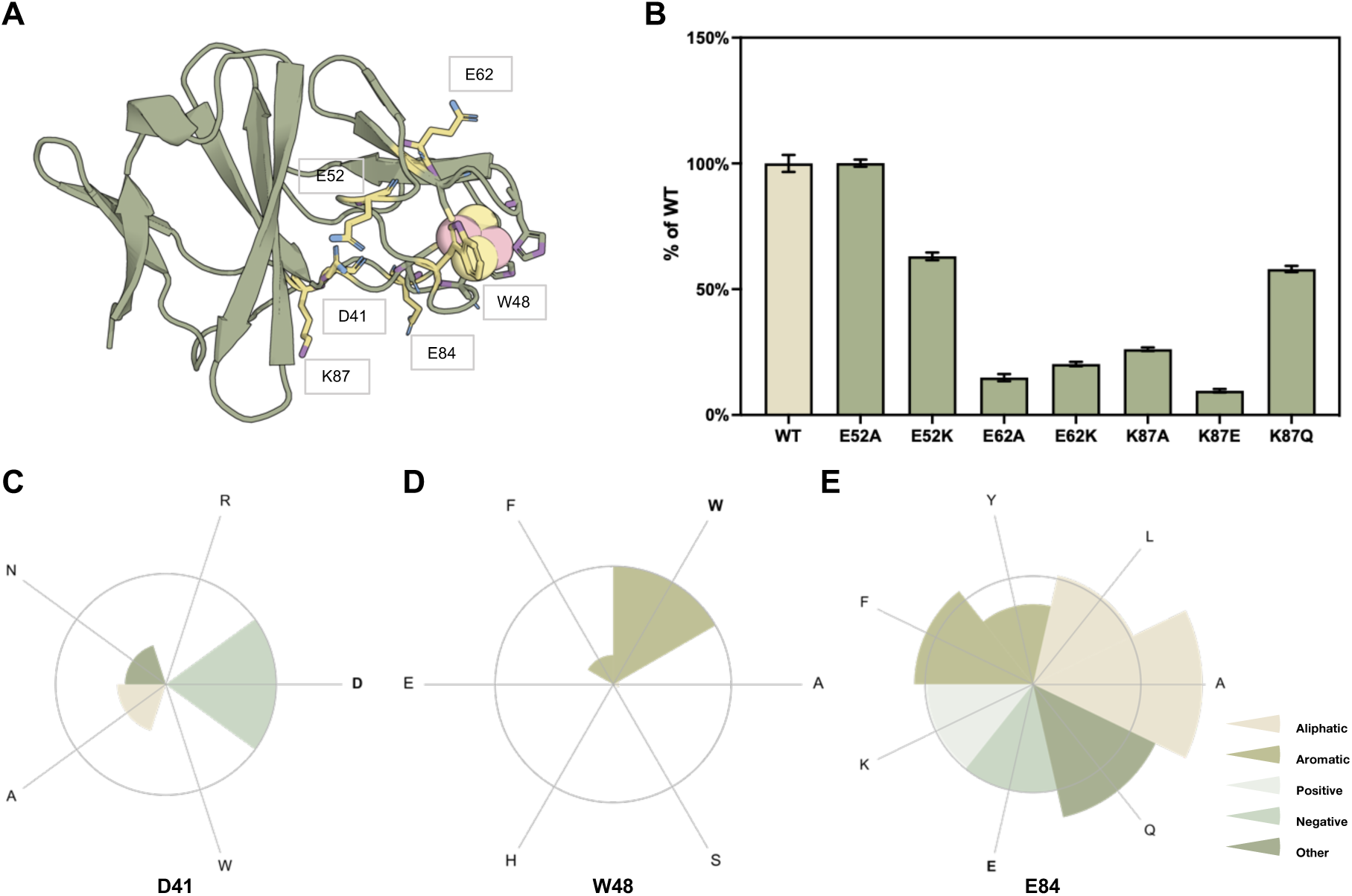
Mutagenesis analysis of critical residues in CDO-Fd. (A) AlphaFold^34^ model of CDO-Fd with critical residues highlighted in yellow. The coordinating residues of Rieske [2Fe–2S] cluster are depicted as sticks. (B–E) Activity comparison of variants at positions E52 (B), E62 (B), K87 (B), D41 (C), W48 (D), and E84 (E). In panel B, wild-type (WT, product concentration normalized to 100%) is displayed as a beige bar, while green bars indicate the relative product concentrations of each variant. In panels C–E, the WT is indicated in bold, and the activity of WT is represented by a grey circle. Colors indicate amino acid physicochemical classes. Reaction conditions: 4 μM Oxy, 30 μM Fd, 10 μM Red, 10 mM **1**, 50 mM D-glucose, 10 U glucose dehydrogenase (GDH), 400 µM NAD^+^, 1 mg mL^-1^ catalase, 1 mM DTT, 50 mM NaP_i_ (pH 7.2), 30°C, 120 rpm, 3 h. Product formation was analyzed by GC-FID. Product concentrations correspond to the combined amounts of **1a** and **1b**. The data points represent the mean, and the error bars represent the standard deviation of duplicate experiments (n = 2).

In NDO-Fd (Figure 3A), substitution of A50 with small or uncharged residues (such as serine or leucine) was well tolerated; however, bulky (tryptophan) or charged (glutamate) substitutions were detrimental to activity (Figure 3B). Previous studies have proposed that two conserved acidic residues (D/E) universally mediate Fd–Oxy binding through electrostatic interactions^21^. However, our results reveal more system-specific behavior. One conserved acidic pair, E52 in CDO-Fd and D54 in NDO-Fd, likely resides at the binding interface; however, the E52A variant showed no significant change, while the D54A variant exhibited increased activity, indicating that this pair does not appear to play a direct or critical role in Fd–Oxy coupling (Figure 2B and 3C). In contrast, residue E62 in CDO-Fd is essential for Oxy interaction. However, the corresponding residue, E63 in NDO-Fd, which also resides at the binding interface, exerts an opposite effect, as removal of an unfavorable electrostatic interaction enhances activity (Figure 3D). Together, these findings suggest that, although these acidic residues are conserved and positioned near the binding interface, their functional roles are not universally preserved across RO families.

**Figure 3.**
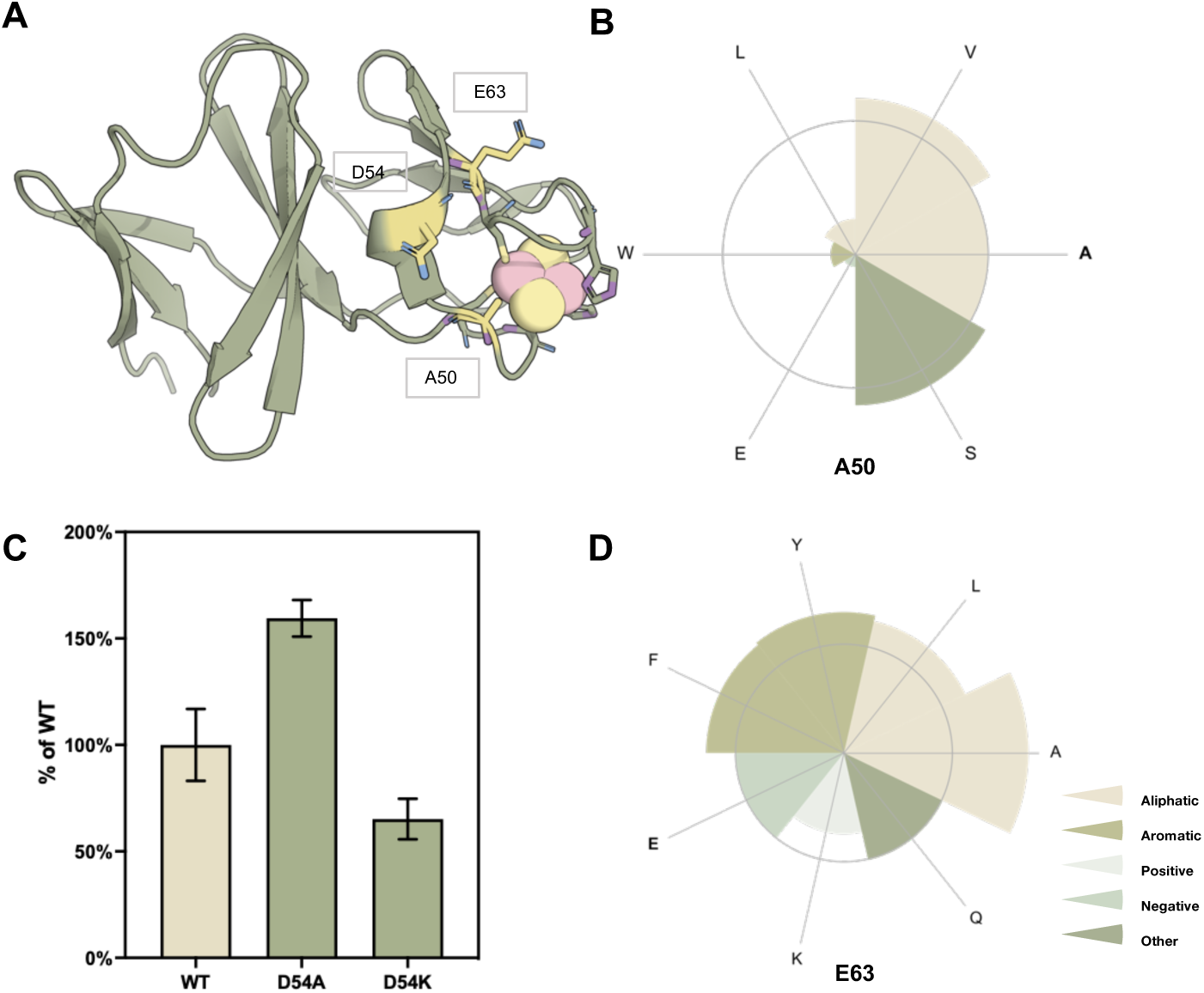
Mutagenesis analysis of critical residues in NDO-Fd. (A) Crystal structure of NDO-Fd (PDB ID: 2QPZ)^39^ with critical residues highlighted in yellow. The coordinating residues of Rieske [2Fe–2S] cluster are depicted as sticks. (B–D) Activity comparison of variants at positions A50 (B), D54 (C) and E63 (D). In panels B and D, the WT is indicated in bold, and the activity of WT is represented by a grey circle. Colors indicate amino acid physicochemical classes. In panel C, WT (product concentration normalized to 100%) is displayed as a beige bar, while green bars indicate the relative product concentrations of each variant. Reaction conditions: 4 μM Oxy, 30 μM Fd, 10 μM Red, 10 mM **1**, 50 mM D-glucose, 10 U glucose dehydrogenase (GDH), 400 µM NAD^+^, 1 mg mL^-1^ catalase, 1 mM DTT, 50 mM NaP_i_ (pH 7.2), 30 °C, 120 rpm, 3 h. Product formation was analyzed by GC-FID. Product concentrations correspond to the combined amounts of **1a** and **1b**. The data points represent the mean, and the error bars represent the standard deviation of duplicate experiments (n = 2).

To pinpoint the Fd regions governing Oxy recognition, we constructed chimeric Fds (Fd-CN_R_C denotes a chimera in which the Rieske cluster-containing segment of CDO-Fd was replaced with the corresponding segment from NDO-Fd; Fd-NC_R_N denotes the construct in which the Rieske cluster-containing segment of NDO-Fd was replaced with that from CDO-Fd) by exchanging the segments containing the Rieske cluster between CDO-Fd and NDO-Fd (Figure 4A and 4B). Additionally, triple mutants targeting unconserved residues (D47, S64 and M67 in CDO-Fd and S49, P65 and Q68 in NDO-Fd) were generated because these positions have different physicochemical properties and are located near the Rieske cluster ligands. Some variants, such as CDO-Fd D47S/S64P/M67Q and the Fd-CN_R_C chimera, exhibited high total protein yields but reduced reddish coloration. This indicates incomplete Rieske [2Fe–2S] cluster incorporation. Thus, the functional concentrations of all Fd variants were recalibrated using the Rieske-specific 325 nm absorption peak with wild-type Fd as the reference (Figure S4). Activity assays using native Reds revealed that CDO-Fd was highly sensitive to minimal sequence perturbations, causing severe loss of activity (Figure 4C). This is likely due to impaired Oxy binding rather than Red mismatch. Conversely, introducing the Rieske region of NDO-Fd into CDO-Fd (Fd-CN_R_C) or constructing the NDO-Fd triple mutant S49D/P65S/Q68M substantially restored ET compatibility with NDO-Oxy (Figure 4C).

**Figure 4.**
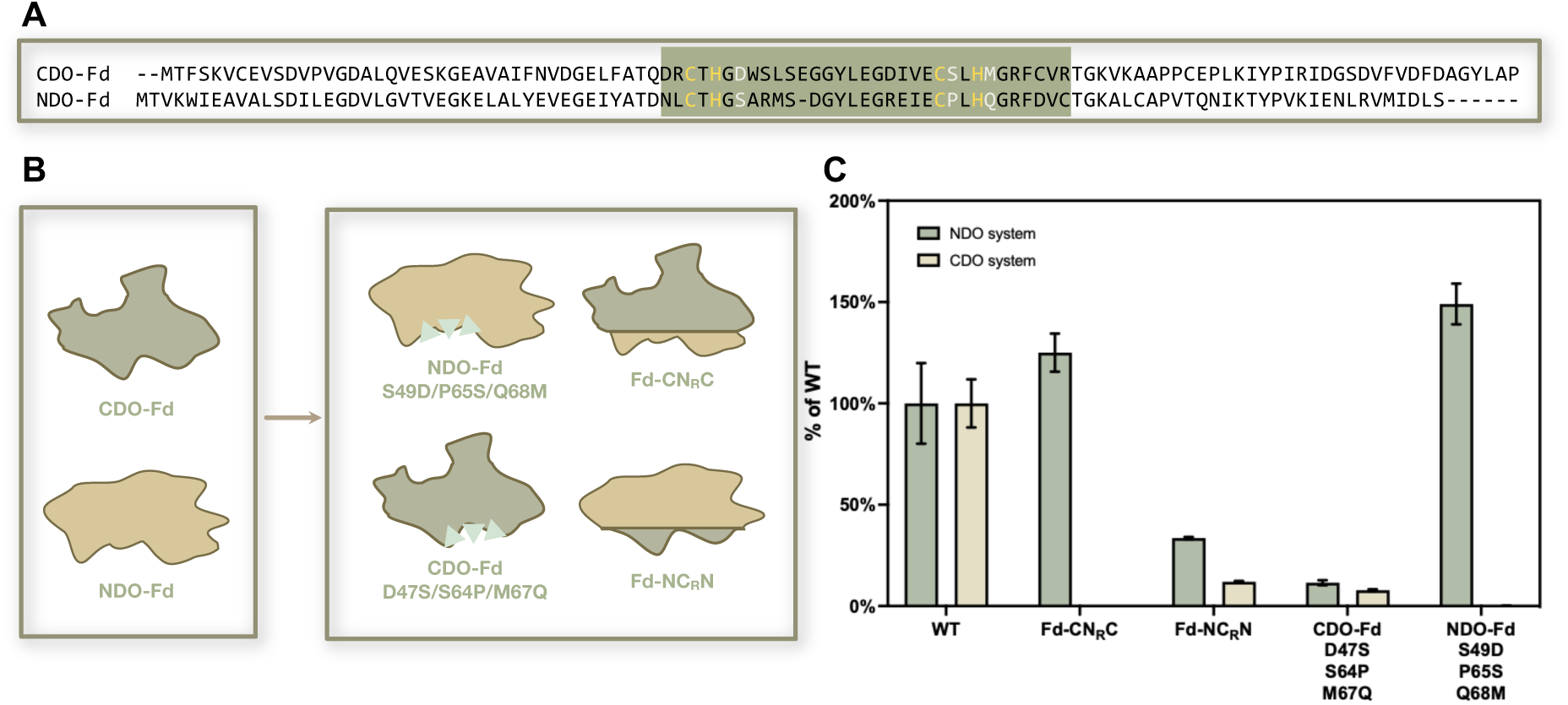
Design and functional evaluation of chimeric ferredoxins. (A) Amino acid sequence alignment of CDO-Fd and NDO-Fd. Residues coordinating the Rieske [2Fe–2S] cluster are marked in yellow. Three unconserved residues selected for mutagenesis are shown in white. Rieske cluster-containing segments exchanged to generate chimeric ferredoxins are highlighted in green. Multiple sequence alignments were performed using Clustal Omega^40^. Default settings were kept for all parameters. (B) Schematic illustration of the chimeric Fds and Fd variants constructed in this study. In Fd-CN_R_C, the Rieske cluster-containing segment of CDO-Fd was replaced with the corresponding segment from NDO-Fd; in Fd-NC_R_N, the Rieske cluster-containing segment of NDO-Fd was replaced with that from CDO-Fd. (C) Assessment of the catalytic activities of the CDO and NDO systems using chimeric Fds and triple-mutant variants. The product concentrations of native systems (WT) were normalized to 100%. The “CDO system” refers to assays performed with CDO-Oxy and CDO-Red, and the “NDO system” refers to assays performed with NDO-Oxy and NDO-Red. Reaction conditions: 4 μM Oxy, 30 μM Fd, 10 μM Red, 10 mM **1**, 50 mM D-glucose, 10 U glucose dehydrogenase (GDH), 400 µM NAD^+^, 1 mg mL^-1^ catalase, 1 mM DTT, 50 mM NaP_i_ (pH 7.2), 30 °C, 120 rpm, 3 h. Product formation was analyzed by GC-FID. Product concentrations correspond to the combined amounts of **1a** and **1b**. The data points represent the mean, and the error bars represent the standard deviation of duplicate experiments (n = 2).

Based on these analyses, A50 in NDO-Fd emerged as a key determinant of redox partner specificity. Examining previously reported sequence alignments across ROs^32^ revealed that this residue position (e.g., W48 in CDO-Fd and A50 in NDO-Fd) is class-dependent: tryptophan predominates in Class IIB Fds, alanine in Class III Fds, and tyrosine in certain plant-type Fds from Class IIA. Encouraged by these results, we constructed a reciprocal NDO–CDO hybrid, combining NDO-Red, NDO-Fd A50W, and CDO-Oxy as separate components. During purification, the A50W variant also exhibited increased total protein yield but reduced coloration, indicating partial Rieske [2Fe–2S] cluster incorporation. After recalibrating the functional concentrations, activity assays showed that this hybrid produced less total product than the native CDO system, yet it markedly outperformed the wild-type NDO–CDO combination (Figure S5). Together, these results demonstrate that the identity of this single residue can reshape redox partner compatibility across RO classes.

### Rational Design of Oxygenase for Enhanced Electron Transfer Efficiency

We hypothesized that a hybrid NDO–CDO system could achieve enhanced ET efficiency through targeted Oxy engineering because the native NDO system exhibited a higher initial reaction rate than the native CDO system (TON_20min_ values of 372 and 66, respectively). Previous structural and simulation analyses identified two potential protein–protein interaction regions within the Oxy: (1) the α-subunit “cap” located at the top of the mushroom-shaped structure^20^, and (2) the α/β-subunit interface situated at the stem.^21^ Guided by the observation that fewer charged residues are critical in NDO-Fd than in CDO-Fd, indicating a reduced dependence on electrostatic interactions, we designed 16 CDO-Oxy variants (Table S6) aimed at minimizing electrostatic and hydrophobic interactions while emulating the characteristics of NDO-Oxy (Figure 5A and 5B). Of these variants, α_E32A, α_W159A and α_D129A exhibited low expression levels and were thus excluded from further analysis. Endpoint activity assays identified three promising variants—α_F144Y, α_A415F, and β_L98K—which exhibited product formation rates 2.6-, 3.1-, and 6.7-fold higher than the CDO-Oxy wild type (Figure 5C). However, combining the three mutations did not yield further improvement (Figure 5D), suggesting possible functional redundancy or structural interference among these sites. Consequently, CDO-Oxy β_L98K was selected as the optimal variant. Time-course assays revealed that the TON_20min_ of this variant increased from 66 in the native CDO system to 205 (Figure 5E), demonstrating a significant enhancement in catalytic efficiency.

**Figure 5.**
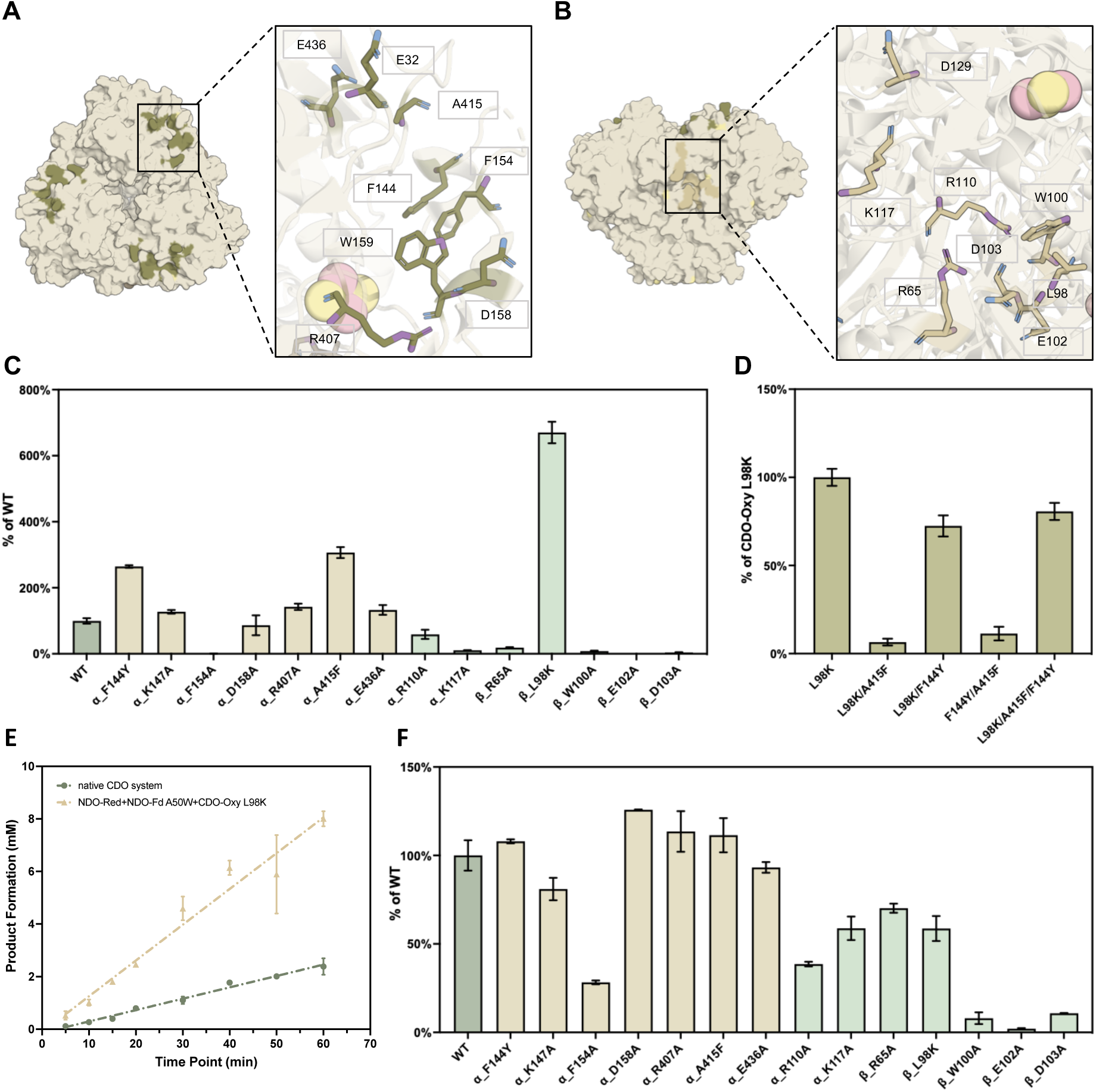
Rational design of CDO-Oxy. (A) Top-surface amino acid residues selected for rational design are shown as green sticks. Residue α_K147 is not shown because it was missing from the crystal structure. (B) Side-surface amino acid residues selected for rational design are shown as beige sticks. In both panels, the left images display the molecular surface of CDO-Oxy (PDB ID: 1WQL)^41^, and the right images show zoomed-in views of the highlighted residues. (C) Activities with NDO-Red and NDO-Fd A50W as redox partner. WT (product concentrations normalized to 100%) is displayed as a grayish green bar. Beige and sage green bars represent the relative product concentrations of variants with amino acid substitutions on the top and side surfaces, respectively. (D) Activities of combinatorial mutants of CDO-Oxy with NDO-Red and NDO-Fd A50W as redox partner. Product concentrations of variant L98K were normalized to 100%, and other variants show relative product concentrations of variants L98K. (E) Time course analysis of total product formation in reactions catalyzed by NDO-Red + NDO-Fd A50W + CDO-Oxy β_L98K compared with the native CDO system. (F) Activities of CDO-Oxy variants with their native redox partner. WT (product concentrations normalized to 100%) is displayed as a grayish green bar. Beige and sage green bars are the relative product concentrations of variants with amino acid substitutions on the top and side surfaces, respectively. Reaction conditions: 4 μM Oxy, 30 μM Fd, 10 μM Red, 10 mM **1**, 50 mM D-glucose, 10 U glucose dehydrogenase (GDH), 400 µM NAD^+^, 1 mg mL^-1^ catalase, 1 mM DTT, 50 mM NaP_i_ (pH 7.2), 30 °C, 120 rpm, 3 h unless indicated otherwise. Product formation was analyzed by GC-FID. Product concentrations correspond to the combined amounts of **1a** and **1b**. The data points represent the mean, and the error bars represent the standard deviation of duplicate experiments (n = 2).

Additionally, all of the designed CDO-Oxy variants were evaluated within the native CDO system (Figure 5F). Notably, variants α_F154A, α_R110A, α_K117A, β_R65A, β_W100A, β_E102A, and β_D103A consistently exhibited reduced product formation in both systems, albeit to varying degrees. Intriguingly, only F154 is located on the top surface, suggesting that there may be multiple binding modes between Fd and Oxy. In contrast, variant β_L98K demonstrated decreased activity exclusively in the native system. These results are consistent with previous findings that residues α_K117 and β_R65 are critical for protein–protein interactions^21^. Specifically, a previous study reported that the β_W100A variant exhibited a 75% decrease in Rieske cluster reduction efficiency relative to the wild type^21^. However, our data show that β_W100A retained only 8% of the wild-type product formation. This implies that β_W100 influences both ET from Fd to Oxy and intramolecular ET from the Rieske cluster to the mononuclear iron center within the Oxy. Although β_W100 is not directly located between these two redox centers in the Oxy, it may play an indirect yet critical role in maintaining the proper protein microenvironment. Since the β_E102A and β_D103A variants nearly abolished activity to a degree comparable to β_W100A, these residues may have similar functional roles in sustaining efficient ET. Nevertheless, the possible involvement of these residues in mediating protein–protein interactions or facilitating ET remains to be verified.

At the same time, our results demonstrate that CDO-Fd E84A enhances product formation and accelerates the initial reaction rate compared to the CDO-Fd wild-type (Figure S6). Therefore, we introduced this mutation into NDO-Fd A50W. The corresponding residue in NDO-Fd is an uncharged glutamine (Q85), and the new variant exhibited improved activity in the engineered hybrid system (NDO-Red, NDO-Fd A50W/Q85A and CDO-Oxy β_L98K). However, the NDO-Fd A50W/E63A double mutant showed a significant decline in product formation (Figure S7), despite E63A’s positive effect in the NDO native system alone. These findings suggest that the beneficial effects of specific Fd positions primarily arise from better interaction with their corresponding Oxy rather than with the Red component. Furthermore, NDO-Fd interacts with CDO-Oxy through a distinct interaction mode compared to its native partner NDO-Oxy.

### Investigating the Redox Partner Recognition and Electron Transfer Mechanisms between Fd and Oxy

According to Marcus’s theory, long-range ET in biological systems is governed by three main factors: the driving force (ΔG), electronic coupling between donor and acceptor (V_DA_), and the reorganization energy (λ)^33^. The driving force primarily depends on the redox gradients along the ET chain. Due to the proximity of tryptophan (W) and alanine (A) to the Rieske cluster in Fd, we first evaluated the redox potentials of the Rieske cluster in both Fd and Oxy. We determined the midpoint redox potentials using electron paramagnetic resonance (EPR) spectroscopy monitored redox titrations. Overall, only minor differences in redox potential were observed among the Fd and Oxy variants (Extended Data Figure 1A−1E). For example, the A50W mutation in NDO-Fd increased the midpoint potential by 15 mV relative to the wild type, whereas the L98K mutation in CDO-Oxy increased the midpoint potential by 8 mV. Notably, the EPR spectra of the Rieske [2Fe–2S]^+^ cluster in the A50W variant revealed subtle differences in the g values, indicating conformational changes in the vicinity of the cluster (Table S7). Specifically, the redox potential difference between NDO-Fd and CDO-Oxy remains within a reasonable range for ET, suggesting that the inability of NDO-Fd to function with CDO-Oxy is unlikely to arise from unfavorable ET thermodynamics only. In addition, the midpoint potential of CDO-Oxy (−159 mV) is 43 mV more positive than that of CDO-Fd and 46 mV more positive than that of NDO-Fd A50W. Furthermore, the midpoint potential of CDO-Oxy β_L98K (−151 mV) is 54 mV more positive than that of NDO-Fd A50W, indicating a slightly larger thermodynamic driving force for electron transfer in this pair.

Electronic coupling is largely dictated by protein–protein affinity, which in turn defines the distance and orientation between donor and acceptor redox centers. To quantify this, we used microscale thermophoresis (MST) to measure binding affinity and determine the dissociation constant (K_D_) between Fd and Oxy. All measured K_D_ values are within the range of 0.73−1.16 μM. The CDO-Oxy β_L98K variant exhibited an affinity for NDO-Fd A50W comparable to that of wild-type CDO-Oxy (Extended Data Figure 1F). Notably, both non-native pairings showed slightly weaker binding than the native CDO-Fd/CDO-Oxy complex, which displayed the lowest K_D_ (Extended Data Figure 1G). However, despite a similar K_D_ to the native complex, the NDO-Fd/CDO-Oxy pairing exhibited poor ET activity, suggesting that binding may trap NDO-Fd in a non-productive orientation. Together, these results suggest that a moderate rather than maximal protein–protein affinity is favorable for efficient ET.

To further elucidate the molecular basis of the Fd–Oxy recognition, we used AlphaFold3^34^ to predict the structures of NDO-Fd A50W and CDO-Oxy β_L98K, and HADDOCK 2.4 web server^35,36^ to model their binding complexes. Docking simulations were performed for the NDO-Fd A50W/CDO-Oxy and NDO-Fd A50W/CDO-Oxy β_L98K complexes. In HADDOCK, docking is typically guided by ambiguous interaction restraints (AIRs), which, in this case, were derived from our mutagenesis data. For the NDO-Fd A50W/CDO-Oxy complex, the interface-active residues were W50 and Q85 of NDO-Fd A50W, and α_K117 and β_R65 of CDO-Oxy; β_L98K was additionally defined as active in the mutant complex. The docking results indicated that Fd binds within a hydrophobic groove at the α/β-subunit interface of Oxy in both complexes but adopts different binding angles in the two Fds (Figure 6A and 6B). Introduction of the β_L98K mutation created a new electrostatic interaction with D54 in NDO-Fd A50W (Figure 6B). While this mutation enhances electrostatic interactions, this gain is likely offset by less favorable desolvation, reduced buried surface area, and weakened van der Waals contacts (Table S8), resulting in little net change in binding affinity. Importantly, formation of this interaction also alters the orientation of NDO-Fd A50W, reducing the distance between the Rieske clusters of Fd and Oxy from 18.2 Å to 16.9 Å. These findings support the notion that targeted charge modulation at the protein interface can strengthen productive complex formation and thereby facilitate ET.

**Figure 6.**
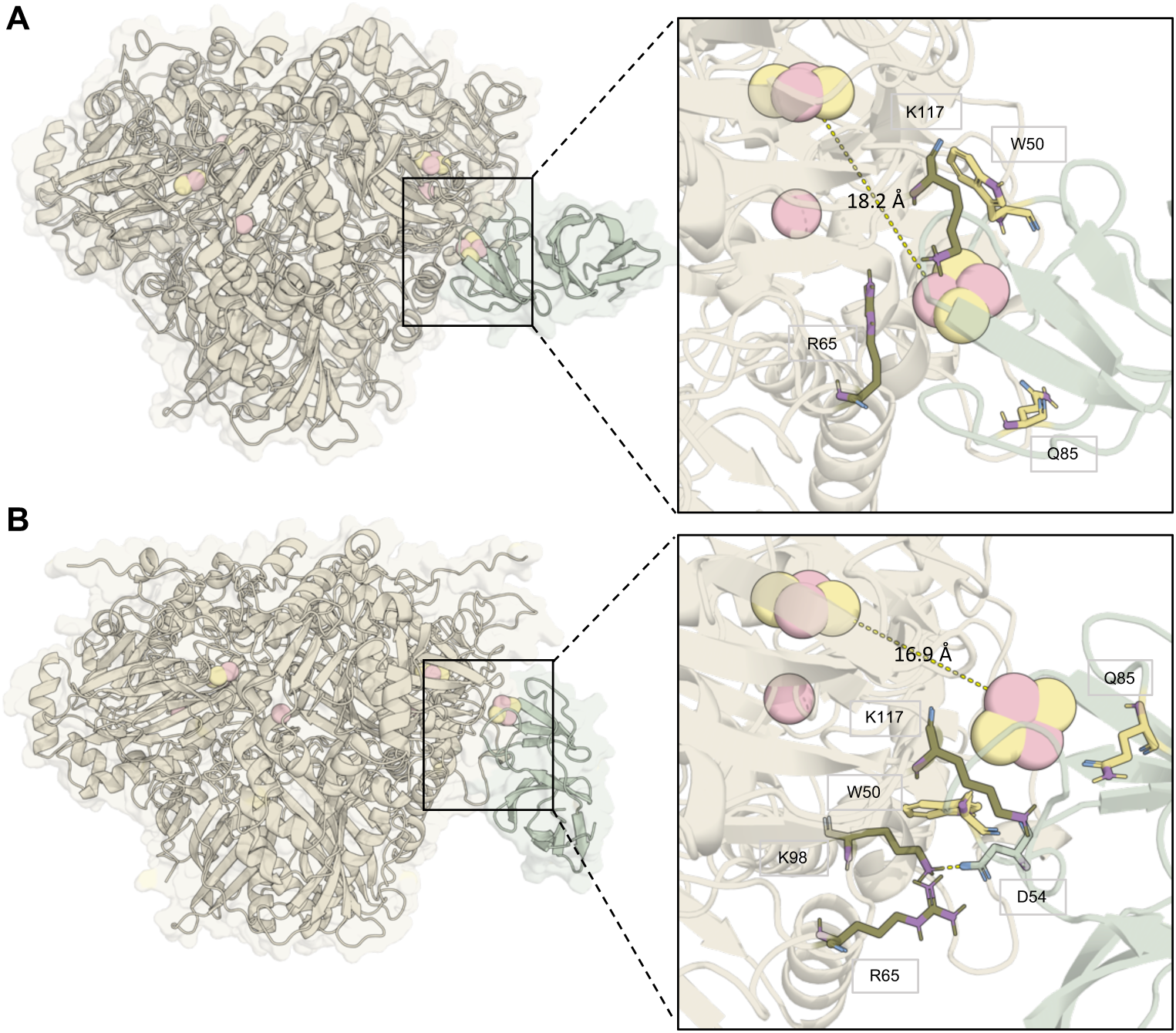
Modelling of ferredoxin–oxygenase binding interfaces. Predicted binding sites of (A) NDO-Fd A50W on CDO-Oxy and (B) NDO-Fd A50W on CDO-Oxy β_L98K. Residues used for docking are shown in green for the oxygenase and yellow for the ferredoxin. In panel B, an intermolecular salt bridge between K98 in the oxygenase and D54 in the ferredoxin was indicated by the yellow dashed line.

Finally, to evaluate the impact of different redox partners and Oxy combinations on unproductive oxygen activation (O₂ uncoupling)^37^, we performed a colorimetric ABTS-HRP assay to quantify the release of H₂O₂ as a reactive oxygen species (ROS) resulting from O₂ uncoupling. However, no significant differences in uncoupling levels were detected between the hybrid systems and the native CDO system (Figure S8).

### Universality of the Redox Partner Recognition Rule in ROs

To evaluate whether the Fd–Oxy recognition principles identified for the NDO and CDO systems extend to other ROs, we selected toluene dioxygenase (TDO) as an additional model system. TDO has been extensively characterized over the past two decades^38^. Like CDO, TDO belongs to Class IIB and shares 55% sequence identity with CDO-Fd, as well as 64% and 51% sequence identity with the α- and β-subunits of CDO-Oxy, respectively. Strikingly, the residue corresponding to W48 in CDO-Fd is also a tryptophan in TDO-Fd (Figure S9A), suggesting that TDO and CDO may follow similar Fd–Oxy recognition principles. Based on these similarities, we hypothesized that the strategy used to reprogram the NDO-Fd and CDO-Oxy interface could also be effective in engineering compatibility between NDO-Fd and TDO-Oxy. Both the wild-type TDO components and the TDO-Oxy β_V99K variant were recombinantly expressed and purified using previously established protocols^30^. The β_V99K mutation in TDO-Oxy was introduced based on sequence alignment, which revealed that position V99 in TDO corresponds to position β_L98 in CDO-Oxy (Figure S9B).

We assessed the catalytic activity of the engineered NDO–TDO hybrid system *in vitro* using toluene **2** as the model substrate (Extended Data Figure 2A). As observed in the CDO system, the TDO-Oxy wild type exhibited low activity when combined with NDO-Fd A50W and was nearly inactive when paired with NDO-Fd (Extended Data Figure 2B). However, the TDO-Oxy β_V99K variant showed significant improvement in product formation in the unnatural system compared to the wild type. Although the engineered NDO–TDO hybrid system did not surpass the catalytic performance of the native TDO system, these results demonstrate that introducing a single mutation at the sequence position equivalent to that identified in the CDO system predictively tunes TDO-Oxy to function with NDO-Red and NDO-Fd A50W as redox partners.

## DISCUSSION

ROs are versatile enzymes that catalyze a variety of oxidative reactions through complex ET processes involving multiple redox partners. Elucidating the molecular determinants of redox partner recognition and compatibility in ROs is crucial for utilizing this class of enzymes in biocatalytic applications. Despite decades of research illuminating critical features common to ROs, the molecular mechanisms governing redox partner recognition and ET pathways in ROs remain surprisingly understudied. In this study, we identified key mechanisms that facilitate productive interactions and ET between Fd and Oxy. We then applied these design principles to predictively tune redox partner recognition in ROs. First, we identified key residues responsible for recognizing redox partners in the CDO and NDO systems through systematic mutagenesis. Notably, the CDO-Fd system exhibited a greater number of charged residues that contribute to protein–protein interactions than the NDO-Fd system. Moreover, the identified alanine and tryptophan residues were found to be conserved within Fds of the same enzyme class. Based on these findings, a hybrid NDO–CDO system was engineered by redesigning the Fd–Oxy interaction interface. This hybrid system displayed markedly enhanced catalytic performance, with the TON_20min_ increasing from 66 to 205. To investigate the interprotein ET mechanism between the engineered NDO-Fd and CDO-Oxy, we conducted a combination of EPR, MST, and computational docking analyses. The results revealed that an optimized redox gradient provides an appropriate driving force for ET in both the native and hybrid system. Meanwhile, finely tuned electrostatic interactions at the protein interface significantly strengthen ET coupling in the hybrid system. Consequently, this refined ET pathway explains the superior catalytic efficiency observed in the engineered NDO–CDO hybrid system. Furthermore, the developed redox partner engineering strategy was successfully extended to the TDO system. In conclusion, the redox partner engineering framework developed in this study provides a rational, generalizable approach to optimizing interprotein ET in multi-component enzyme systems. Our findings deepen the mechanistic understanding of ET in ROs and establish fundamental design principles for predictively tuning hybrid or surrogate electron transport chains. Thus, this study paves the way for developing versatile, highly efficient biocatalytic systems based on ROs.

## METHODS

### Site-directed mutagenesis

Site-directed mutagenesis was performed to introduce specific nucleotide substitutions into target plasmids. Mutagenic primers carrying the desired base changes were designed according to the QuikChange protocol^42^. The 25 µL reactions were composed of 12.5 µL Q5 High-Fidelity 2X Master Mix, 1 ng µL^-1^ plasmid template and 1 µM primer. Reactions were run using the following program: 98 °C 30 s, (98 °C 15 s, 65 °C 20 s and 72 °C 3 min 45 s) for 26 cycles, and 72 °C for 2 min. Following amplification, the parental (template) DNA was digested with 1 µL DpnI (37 °C, 1 h). The resulting 5 µL PCR product was directly used for transformation of *E. coli* DH5α competent cells via heat shock. Positive clones were verified by DNA sequencing (Macrogen Europe, the Netherlands). Primer sequences used for site-directed mutagenesis and a detailed list of CDO-Oxy variants generated in this study can be found in the Supplementary Information (Table S4, Table S6).

### Expression and purification of recombinant proteins

Heterologous expression of glucose dehydrogenase from *Bacillus megaterium* (GDH) was performed using *E. coli* BL21(DE3) as the expression host^43^. Protein expression and purification were carried out following the same general procedures described below for the Rieske oxygenase (RO) protein components, with the only difference that culture medium was supplemented with 100 µg mL⁻¹ ampicillin. Samples were frozen in liquid nitrogen for storage. Heterologous expression of RO protein components was performed using *E. coli* JM109(DE3) as the expression host. High-cell-density precultures were prepared from freshly transformed *E. coli* JM109(DE3) colonies grown on LB agar plates supplemented with 50 µg mL⁻¹ kanamycin. Single colonies were used to inoculate baffled Erlenmeyer flasks containing *lysogeny* broth (LB) medium supplemented with 50 µg mL⁻¹ kanamycin. The inoculated precultures were incubated overnight at 37 °C and 180 rpm, and subsequently used to inoculate the main expression cultures. Main cultures were grown in baffled Erlenmeyer flasks containing terrific broth (TB) medium supplemented with 50 µg mL⁻¹ kanamycin, and inoculated to an initial OD_600_ of 0.1. Cultures were incubated at 37 °C and 120 rpm until reaching an OD_600_ of approximately 1.0, at which point protein expression was induced by the addition of 100 µM IPTG. After induction, the cultures were incubated for 16–19 h at 20 °C and 120 rpm. Cell harvesting, washing, and lysis were performed according to a previously optimized purification protocol^30^. The clarified lysate was incubated with 2 mL Ni-sepharose resin (1 column volume (CV) = 2 mL) for 1 h, followed by washing with 5 CV of washing buffer (50 mM NaP_i_, 40 mM imidazole, 300 mM NaCl, 10% glycerol, pH 7.2). Elution was carried out using elution buffer (50 mM NaP_i_, 400 mM imidazole, 300 mM NaCl, 10% glycerol, pH 7.2). Elution fractions corresponding to 0.5–2×CV were collected for downstream processing. To exchange the buffer to the final storage composition (50 mM NaP_i_, 300 mM NaCl, 10% glycerol, pH 7.2), purified proteins were passed through PD-10 desalting columns (Cytiva). Final protein concentrations were determined using a colorimetric Coomassie protein assay, as described previously^30^. All samples were freshly prepared and immediately used for subsequent experiments.

### *In vitro* biochemical assays and product analysis

*In vitro* reactions were prepared in 20 mL tightly-sealed glass vials on a 1 mL scale. Unless stated otherwise, a standard reaction solution contained purified protein components at concentrations of 4 μM oxygenase (Oxy), 30 μM ferredoxin (Fd) and 10 μM reductase (Red). In addition, standard reactions contained 50 mM D-glucose, 10 U mL^−1^ GDH, 1 mM dithiothreitol (DTT), 1 mg mL^−1^ catalase, and 400 µM NAD^+^. Substrate was added directly to the reaction solutions prepared in 50 mM NaP_i_ (pH 7.2) to a final concentration of 10 mM. For toluene **2**, 500 mM stock solutions were prepared in DMSO. Reactions were carried out in an incubation shaker at 30 °C at 120 rpm for the indicated reaction times. Organic compounds in the aqueous mixtures were extracted using dichloromethane (DCM) containing 2 mM acetophenone as an internal standard (IS). Extraction efficiency was improved by saturating the aqueous phase with sodium chloride (NaCl). Organic extracts were dried over anhydrous MgSO_4_ before analysis by gas chromatography (GC, see Supplementary information). The conversions of analytical scale reactions are calculated based on the integrals of the GC-FID analysis.

### Redox titration of Fd and Oxy

The titration was performed within an anaerobic chamber (Coy Laboratory Products) at room temperature. To maintain anaerobic conditions, oxygen was removed from the buffer solutions (50 mM NaP_i_, 300 mM NaCl, 10% glycerol, pH 7.2) using nitrogen displacement. The following dyes were solved with anaerobic buffer to a stock concentration of 160 μM: *N*,*N*,*N*’,*N*’-tetramethyl-*p*-phenylenediamine, 2,6-dichlorophenol indophenol, phenazine ethosulfate, methylene blue, resorufine, indigodisulfonate, 2-hydroxy-1,4-naphtaquinone, anthraquinone-2-sulfonate, phenosafranin, safranin O, neutral red, benzyl viologen, and methyl viologen. The final concentration of each mediator was stoichiometric or higher than the Rieske cluster concentration. Samples were first reductively titrated with sodium dithionite to a potential of circa −400 mV, followed by oxidative titration with potassium ferricyanide. After equilibration at the desired potential, a 0.2 mL sample was anaerobically transferred to an electron paramagnetic resonance (EPR) tube and immediately frozen in liquid nitrogen. The titration was performed in oxidative direction after initial reduction of the titration mixture to a potential of circa −400 mV. Potentials were measured with a platinum electrode and an Ag/AgCl reference electrode. All reported values are with respect to the normal hydrogen electrode (NHE). A detailed protocol for the EPR monitored redox titration has been previously published^44^.

### Electron paramagnetic resonance measurement

EPR spectra were acquired using a Bruker EMXplus spectrometer equipped with a helium-flow cryostat operating at a temperature of 19 K. The experimental parameters were set as follows: microwave frequency 9.406 GHz, microwave power 0.63 mW, a modulation frequency of 100 kHz, and a modulation amplitude of 10 G. Data analysis was performed as previously described^44^.

### Microscale thermophoresis assay

Microscale thermophoresis (MST) was used to determine ligand–target binding affinities by monitoring binding-induced changes in molecular size, charge, or solvation^45^. CDO-Oxy WT and CDO-Oxy L98K were fluorescently labeled using an NHS Labeling Kit (NanoTemper Technologies) and used as target proteins at 50 nM. Measurements were performed in PBS-T binding buffer (PBS containing 0.05% Tween) on a MONOLITH NT.115 instrument (NanoTemper Technologies) at 25 °C. For each assay, 10 µL of labeled target protein was mixed with 10 µL of a 16-point 2-fold serial dilution of ligand protein at room temperature. CDO-Fd WT and NDO-Fd WT were assayed from a starting concentration of 25 µM, whereas NDO-Fd A50W was assayed from a starting concentration of 50 µM. Data were analyzed using MO.Affinity Analysis Software. Measurements were performed in duplicate. During analysis, capillary scans and thermophoresis traces were evaluated for assay quality, and outliers were excluded where appropriate if they showed signs of aggregation or abnormal fluorescence behavior. The single-experiment dataset reports used to generate the final fitted curves are provided in the Supplementary Information.

### Computational methods

Protein–protein docking was performed using the HADDOCK 2.4 web server^36^. The three-dimensional structures of CDO-Fd, NDO-Fd A50W, and CDO-Oxy β_L98K were modeled using AlphaFold3^34^. AlphaFold predictions were obtained using default settings, with full-length amino acid sequences provided as input and no structural templates specified. For each protein, the top-ranked model based on the AlphaFold confidence metrics was selected for subsequent analyses. The crystal structure of wild-type CDO-Oxy was obtained from the Protein Data Bank (PDB ID: 1WQL)^41^. Prior to docking, protein structures were prepared using PyMOL, including removal of solvent molecules and heteroatoms. Docking calculations were performed using the default HADDOCK parameters unless stated otherwise. Statistical analyses of the Fd–Oxy docking results can be found in the Supplementary Information (Table S8). The best-ranked clusters were selected for further analysis.

## Data availability

Data relating to the materials and methods, experimental procedures, mechanistic studies, standard curves, gas chromatography spectra, EPR spectra and protein sequences are available in the Supplementary Information. All raw data are available from the corresponding author upon request. Protein accession codes: Q51747, Q51746, Q51744, Q51743, Q52126, P0A185, P0A110, P0A112, A5W4E9, A5W4F0, A5W4F2, A5W4F1. The crystal structure data used in this study are available in the Protein Data Bank under accession codes 1WQL and 2QPZ.

## Acknowledgements

Hui Miao was supported by a PhD scholarship from the China Scholarship Council (CSC No. 202309110011). PLH acknowledges support from COST Action FeSImmChemNet (CA21115) supported by COST (European Cooperation in Science and Technology) and funding from the European Union’s Horizon Europe research and innovation programme under grant agreement no. 101183014 (project McGEA).

## Author Contributions

H.M. and P.-L.H. designed and performed most of the experiments and data evaluation. R.O. contributed knowledge about protein biochemistry and binding affinity experiments. H.M. and S.S. conceived the project and S.S. directed it. H.M. wrote the manuscript and all authors jointly edited the manuscript. All authors approved the final version of the manuscript.

### Competing interests

The authors declare no competing interests.

**Correspondence and requests for materials** should be addressed to Sandy Schmidt.

## EXTENDED DATA SETS

**Extended Data Figure 1.**
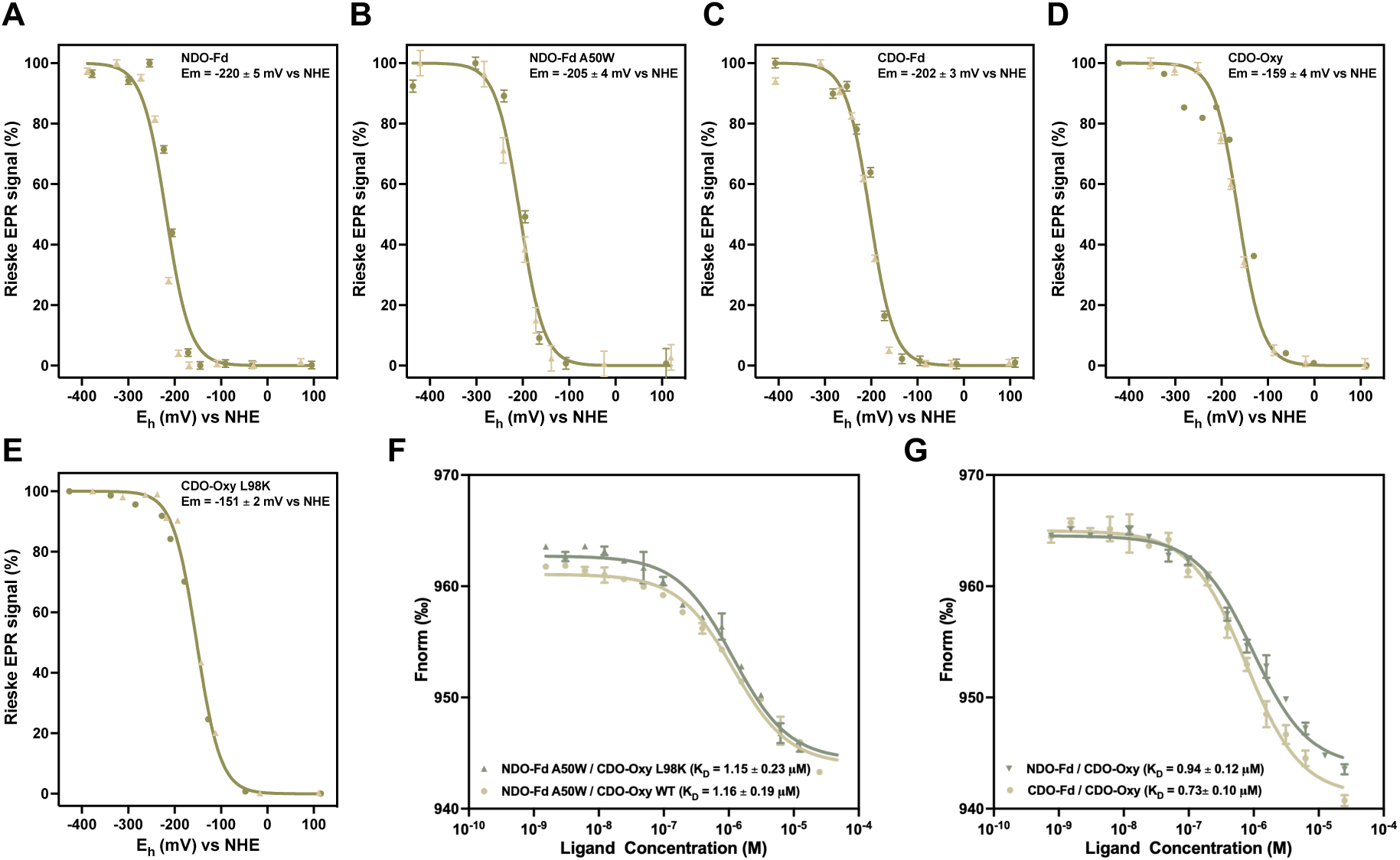
Mechanistic investigation of redox partner recognition and electron transfer. EPR redox titration curves of the Rieske cluster for (A) NDO-Fd, (B) NDO-Fd A50W, (C) CDO-Fd, (D) CDO-Oxy, and (E) CDO-Oxy β_L98K. For each sample, titrations were performed in duplicate and are shown in beige and green, respectively. Error bars represent noise in the EPR spectra. (F) MST binding assay between ligand NDO-Fd A50W and target CDO-Oxy WT or L98K. (G) MST binding assay between ligand CDO-Fd or NDO-Fd and target CDO-Oxy WT. Error bars represent residuals between observed data and fitted values obtained from global K_D_ fitting.

**Extended Data Figure 2.**
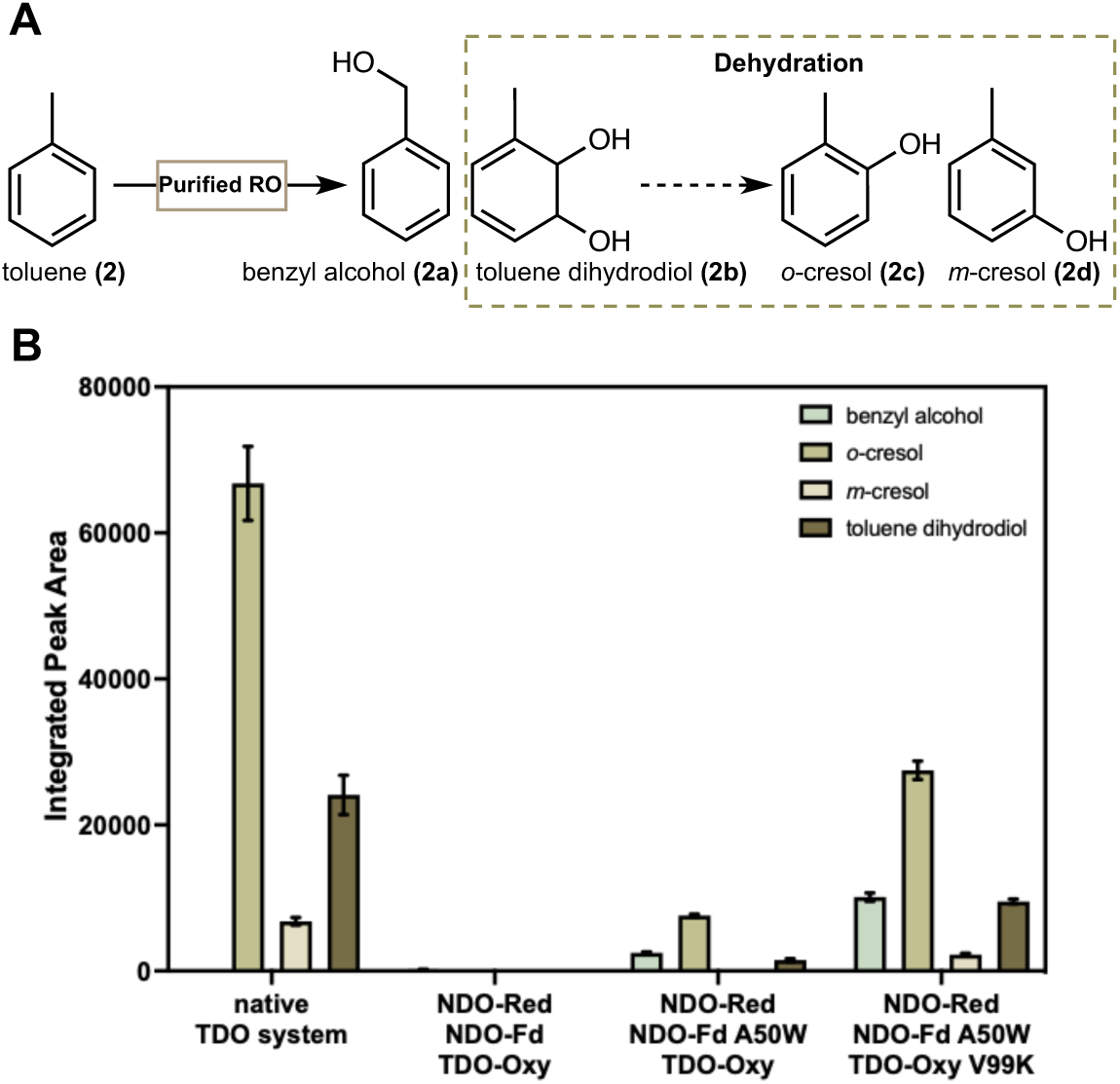
Evaluation of electron transfer compatibility in the NDO–TDO hybrid system. (A) *In vitro* conversion of **2** to **2a** and **2b** catalyzed by TDO; **2c** and **2d** arise from dehydration of **2b**. (B) Assessment of the activity of NDO–TDO hybrid system. Reaction conditions: 4 μM Oxy, 30 μM Fd, 10 μM Red, 10 mM **2**, 50 mM D-glucose, 10 U glucose dehydrogenase (GDH), 400 µM NAD^+^, 1 mg mL^-1^ catalase, 1 mM DTT, 50 mM NaP_i_ (pH 7.2), 30 °C, 120 rpm, 3 h. Samples were analyzed by GC-FID by integrating the peak areas corresponding to **2a**, **2b**, **2c** and **2d**. The data points represent the mean, and the error bars represent the standard deviation of duplicate experiments (n = 2).

## REFERENCES

1. Coulter, E. D. & Ballou, D. P. Non-haem iron-containing oxygenases involved in the microbial biodegradation of aromatic hydrocarbons. Essays Biochem. 34, 31–49 (1999).

2. Yoshiyama-Yanagawa, T. et al. The conserved Rieske oxygenase DAF-36/Neverland is a novel cholesterol-metabolizing enzyme. J. Biol. Chem. 286, 25756–25762 (2011).

3. Gray, J. et al. A small family of LLS1-related non-heme oxygenases in plants with an origin amongst oxygenic photosynthesizers. Plant Mol. Biol. 54, 39–54 (2004).

4. Perry, C., De Los Santos, E. L. C., Alkhalaf, L. M. & Challis, G. L. Rieske non-heme iron-dependent oxygenases catalyse diverse reactions in natural product biosynthesis. Nat. Prod. Rep. 35, 622–632 (2018).

5. Barry, S. M. & Challis, G. L. Mechanism and catalytic diversity of Rieske non-heme iron-dependent oxygenases. ACS Catal. 3, 2362–2370 (2013).

6. Runda, M. E., de Kok, N. A. W. & Schmidt, S. Rieske oxygenases and other ferredoxin-dependent enzymes: Electron transfer principles and catalytic capabilities. ChemBioChem 24, e202300078 (2023).

7. Lee, J. & Zhao, H. Mechanistic studies on the conversion of arylamines into arylnitro compounds by aminopyrrolnitrin oxygenase: Identification of intermediates and kinetic studies. Angew. Chem. Int. Ed. 45, 622–625 (2006).

8. Summers, R. M., Louie, T. M., Yu, C. L. & Subramanian, M. Characterization of a broad-specificity non-haem iron *N*-demethylase from *Pseudomonas putida* CBB5 capable of utilizing several purine alkaloids as sole carbon and nitrogen source. Microbiology (N Y). 157, 583–592 (2011).

9. Priefert, H., Rabenhorst, J. & Steinbüchel, A. Molecular characterization of genes of *Pseudomonas* sp. Strain HR199 involved in bioconversion of vanillin to protocatechuate. J. Bacteriol. 179, 2595–2607 (1997).

10. D’Ordine, R. L. et al. Dicamba monooxygenase: Structural insights into a dynamic Rieske oxygenase that catalyzes an exocyclic monooxygenation. J. Mol. Biol. 392, 481–497 (2009).

11. Kancharla, P., Lu, W., Salem, S. M., Kelly, J. X. & Reynolds, K. A. Stereospecific synthesis of 23-hydroxyundecylprodiginines and analogues and conversion to antimalarial premarineosins via a Rieske oxygenase catalyzed bicyclization. J. Org. Chem. 79, 11674–11689 (2014).

12. Piskol, F. et al. Two-component carnitine monooxygenase from *Escherichia coli*: functional characterization, inhibition and mutagenesis of the molecular interface. Biosci. Rep. 42, (2022).

13. Ferraro, D. J., Gakhar, L. & Ramaswamy, S. Rieske business: Structure-function of Rieske non-heme oxygenases. Biochem. Biophys. Res. Commun. 338, 175–190 (2005).

14. Runda, M. E., Miao, H., de Kok, N. A. W. & Schmidt, S. Developing hybrid systems to address oxygen uncoupling in multi-component Rieske oxygenases. J. Biotechnol. 389, 22–29 (2024).

15. Lukowski, A. L., Liu, J., Bridwell-Rabb, J. & Narayan, A. R. H. Structural basis for divergent C–H hydroxylation selectivity in two Rieske oxygenases. Nat. Commun. 11, 2991 (2020).

16. Tian, J. et al. Custom tuning of Rieske oxygenase reactivity. Nat. Commun. 14, 5858 (2023).

17. Zhu, Z. et al. Development of engineered ferredoxin reductase systems for the efficient hydroxylation of steroidal substrates. ACS Sustainable Chem. Eng. 8, 16720–16730 (2020).

18. Nojiri, H. Structural and molecular genetic analyses of the bacterial carbazole degradation system. *Biosci., Biotechnol.*, Biochem. 76, 1–18 (2012).

19. Inoue, K. et al. Specific interactions between the ferredoxin and terminal Oxygenase components of a class IIB Rieske nonheme iron oxygenase, carbazole 1,9a-dioxygenase. J. Mol. Biol. 392, 436–451 (2009).

20. Ashikawa, Y. et al. Electron transfer complex formation between oxygenase and ferredoxin components in Rieske nonheme iron oxygenase system. Structure 14, 1779–1789 (2006).

21. Tsai, P. C. et al. The α- and β-subunit boundary at the stem of the mushroom- and α3β3-type oxygenase component of Rieske non-heme iron oxygenases is the Rieske-type ferredoxin-binding site. Appl. Environ. Microbiol. 88, e00835–22 (2022).

22. Wolfe, M. D. & Lipscomb, J. D. Hydrogen peroxide-coupled *cis*-diol formation catalyzed by naphthalene 1,2-dioxygenase. J. Biol. Chem. 278, 829–835 (2003).

23. Neibergall, M. B., Stubna, A., Mekmouche, Y., Münck, E. & Lipscomb, J. D. Hydrogen peroxide dependent *cis*-dihydroxylation of benzoate by fully oxidized benzoate 1,2-dioxygenase. Biochemistry 46, 8004–8016 (2007).

24. He, J., Liu, X. & Li, C. Engineering electron transfer pathway of cytochrome P450s. Molecules 29, 2480 (2024).

25. Li, S., Du, L. & Bernhardt, R. Redox partners: Function modulators of bacterial P450 enzymes. Trends Microbiol. 28, 445–454 (2020).

26. Meng, S. et al. Introduction of aromatic amino acids in electron transfer pathways yielded improved catalytic performance of cytochrome P450s. Chin. J. Catal. 49, 81–90 (2023).

27. Yu, J., Ge, J., Yu, H. & Ye, L. Improved bioproduction of the nylon 12 monomer by combining the directed evolution of P450 and enhancing Heme synthesis. Molecules 28, 1758 (2023).

28. Sagadin, T., Riehm, J., Putkaradze, N., Hutter, M. C. & Bernhardt, R. Novel approach to improve progesterone hydroxylation selectivity by CYP106A2 via rational design of adrenodoxin binding. FEBS J. 286, 1240–1249 (2019).

29. Sun, C., et al. Establishing an efficient electron transfer system for P450 enzyme OleP to improve the biosynthesis of murideoxycholic acid by redox partner engineering. Angew. Chem., Int. Ed. 64, e202423209 (2025).

30. Runda, M. E., Kremser, B., Özgen, F. F. & Schmidt, S. An optimized system for the study of Rieske oxygenase-catalyzed hydroxylation reactions in vitro. ChemCatChem 15, e202300371 (2023).

31. Leggate, E. J. & Hirst, J. Roles of the disulfide bond and adjacent residues in determining the reduction potentials and stabilities of respiratory-type Rieske clusters. Biochemistry 44, 7048–7058 (2005).

32. Nam, J. W. et al. Crystal structure of the ferredoxin component of carbazole 1,9a-dioxygenase of *Pseudomonas resinovorans* strain CA10, a novel Rieske non-heme iron oxygenase system. Proteins: Struct., Funct., Bioinf. 58, 779–789 (2005).

33. Page, C. C., Moser, C. C., Chen, X. & Dutton, P. L. Natural engineering principles of electron tunnelling in biological oxidation-reduction. Nature 402, 47–52 (1999).

34. Abramson, J. et al. Accurate structure prediction of biomolecular interactions with AlphaFold 3. Nature 630, 493–500 (2024).

35. Honorato, R. V. et al. Structural biology in the clouds: The WeNMR-EOSC ecosystem. Front. Mol. Biosci. 8, 729513 (2021).

36. Honorato, R. V. et al. The HADDOCK2.4 web server for integrative modeling of biomolecular complexes. Nat. Protoc. 19, 3219–3241 (2024).

37. Cano, A. & Arnao, M. B. An end-point method for estimation of the total antioxidant activity in plant material. Phytochem. Anal. 9, 196–202 (1998).

38. Friemann, R. et al. Structures of the multicomponent Rieske non-heme iron toluene 2,3-dioxygenase enzyme system. *Acta Crystallogr.*, Sect. D: Biol. Crystallogr. 65, 24–33 (2009).

39. Brown, E. N. et al. Determining Rieske cluster reduction potentials. J. Biol. Inorg. Chem. 13, 1301–1313 (2008).

40. Madeira, F. et al. The EMBL-EBI job dispatcher sequence analysis tools framework in 2024. Nucleic Acids Res. 52, W521–W525 (2024).

41. Dong, X. et al. Crystal structure of the terminal oxygenase component of cumene dioxygenase from *Pseudomonas fluorescens* IP01. J. Bacteriol. 187, 2483–2490 (2005).

## REFERENCES (METHODS)

42. Liu, H. & Naismith, J. H. An efficient one-step site-directed deletion, insertion, single and multiple-site plasmid mutagenesis protocol. BMC Biotechnol. 8, 91 (2008).

43. Prats Luján, A., Bhat, M. F., Saravanan, T. & Poelarends, G. J. Chemo- and enantioselective photoenzymatic ketone reductions using a promiscuous flavin-dependent nitroreductase. ChemCatChem 14, e202200043 (2022).

44. Hagedoorn, P. L., van der Weel, L. & Hagen, W. R. EPR monitored redox titration of the cofactors of Saccharomyces cerevisiae Nar1. J. Visualized Exp. 51611 (2014).

45. Wienken, C. J., Baaske, P., Rothbauer, U., Braun, D. & Duhr, S. Protein-binding assays in biological liquids using microscale thermophoresis. Nat. Commun. 1, 100 (2010).

