## Supplementary Information for "Decoding and Reprogramming Redox Partner Specificity in Rieske Oxygenases for Enhanced Catalytic Activity"

### 1. Supporting Methods

#### 1.1 General materials

Unless stated otherwise, all chemicals were purchased from commercial sources, such as Sigma-Aldrich, Merck, TCI and BLDpharm in the highest purity available and used without any pre-treatment. Solvents dichloromethane (DCM) and dimethyl sulfoxide (DMSO) were used in HPLC grade. Enzymes for Gibson assembly and site-directed mutagenesis were purchased from New England Biolabs. Catalase (from bovine liver), thrombin (from bovine plasma), peroxidase (from horseradish), and 2,2'-azino-bis(3-ethylbenzothiazoline-6-sulfonic acid) diammonium salt (ABTS) were purchased form Sigma-Aldrich. Kits for PCR product purification, DNA gel extraction, and plasmid isolation were obtained from Qiagen. The service of Macrogen Europe (the Netherlands) was used for DNA sequencing. Synthetic genes were purchased from TWIST Bioscience. Plasmids encoding the CDO and NDO systems originated from our previous work[1,2]. Plasmid encoding the GDH was kindly provided by Prof. Gerrit J. Poelarends[3]. GC-FID analysis was performed on a Shimadzu GC-2010 Plus equipped with an AOC-20i auto-injector. GC–MS analysis was performed on a Shimadzu GCMS-QP2010 SE system equipped with an AOC-20i auto-injector module. Column specifications and conditions for analyses are described in Section 1.3 and 1.4.

#### 1.2 Plasmid constructs

Plasmid constructs for the expression of TDO components were generated using Gibson assembly. Synthetic genes for each component were purchased from TWIST Bioscience and codon-optimized using the company's algorithm. DNA fragments required for assembly were prepared in linear form with appropriate Gibson overhangs. Specifically, both the vector backbone, pET-28a(+), and the insert fragment were amplified by PCR. To remove residual template DNA, PCR products were treated with DpnI (37 °C, 1 h) and subsequently purified. Purified PCR products were directly used for Gibson assembly. The reaction was set up using a vector-to-insert molar ratio of 1:5. Calculated DNA amounts (total volume of 10 µL) were combined with 15 µL of Gibson master mix. After incubation at 50 °C for 1 h, 4 µL of the assembly mixture was used for heat-shock transformation of *E. coli* DH5α competent cells. Transformants carrying the correct construct were identified by colony PCR and verified by DNA sequencing. Oligonucleotides used for DNA amplification are listed in Table S4.

#### 1.3 Gas chromatography (GC) analysis

Analysis was performed with a GC-FID (GC-2010, Shimadzu, Kyoto, Japan) equipped with an OPTIMA 5MS column (30.0 m x 0.25 mm, 0.25 µm film thickness, Macherey-Nagel, Düren, Germany). Detailed GC–FID conditions used for the separation and quantification of model substrates and their hydroxylated products from *in vitro* reactions are summarized in Table S1 and Table S2. A representative GC–FID chromatogram is shown in Figure S14 and Figure S15.

The absolute concentrations of indanone were determined using a calibration curve established over a concentration range of 0–5 mM (Figure S11).

For *cis*-1,2-indandiol (**1b**), quantification was based on a racemic indandiol standard. Since the racemate contains equimolar amounts of *cis*- and *trans*-1,2-indandiol, the effective concentration of *cis*-1,2-indandiol (**1b**) used for calibration was assumed to be one-half of the total indandiol concentration. Calibration was conducted over a total indandiol concentration range of 0–10 mM (Figure S12).

1*H*-indenol (**1a**) was not commercially available and its synthesis is challenging due to the risk of the formation of explosive hydroperoxide intermediates (indene-1-peroxide)[4]. Because of that, the calibration of indanone (Figure S11) was used to determine the absolute concentration of 1*H*-indenol (**1a**) via approximation. Therefore, a relative response factor (RRF) was determined based on the effective carbon numbers (ECNs) of 1*H*-indenol (**1a**) and indanone as the reference compound. The related ECNs were evaluated based on the calculation model of Jorgensen et al.[5], and are summarized in Table S5.

The response factors (RFs) listed in Table S5 were calculated using **Equation 1**.

| | |
|---|---|
| $RF\ compound = \frac{ECN\ (compound)}{\mathrm{ECN\ (reference)}}$ | **Equation 1** |

The ratio between the two RFs results in the RRF (**Equation 2**).

| | |
|---|---|
| $RFF = \frac{\mathrm{RF(1}H\mathrm{indenol)}}{\mathrm{RF(indanone)}} = 1.03$ | **Equation 2** |

Finally, the concentration of 1*H*-indenol was calculated by using **Equation 3**.

| | |
|---|---|
| $\mathrm{c(indenol)} = \frac{\mathrm{IPA(indenol)}}{\mathrm{IPA(indanone)}} \times \frac{1}{\mathrm{RFF}} \times c\mathrm{(indanone)}$ | **Equation 3** |

Based on the linear regression for indanone (Figure S11) in combination with Equation 3, an approximated ratio between the concentration of 1*H*-indenol (**1a**) and the integrated peak area (IPA) could be established, corresponding to **Equation 4.**

| | |
|---|---|
| $\mathrm{c(indenol)} = \frac{\mathrm{IPA(indenol)}}{\mathrm{IPA(}IS\mathrm{)}} \times 1.60052$ | **Equation 4** |

This approximation was used for the quantification of 1H-indenol (**1a**) throughout this study.

#### 1.4 GC–MS analysis

Analysis was performed with a GCMS-QP2010 SE system (Shimadzu, Kyoto, Japan) equipped with an OPTIMA 5MS column (30.0 m x 0.25 mm, 0.25 µm film thickness, Macherey-Nagel, Düren, Germany). Detailed GC–MS conditions used for the detection of toluene and corresponding hydroxylated products from *in vitro* reactions are summarized in Table S3. Representative GC–MS chromatograms are shown in Figures S16–S20. Compound identification was based on comparison of the obtained mass spectra with reference databases and previously reported literature data[6].

##### 1.5 Quantification of $H_2O_2$ using colorimetric ABTS-HRP assay

The concentration of hydrogen peroxide ($H_2O_2$) in aqueous reaction samples was determined using a colorimetric assay based on the HRP-catalyzed oxidation of ABTS. In the presence of $H_2O_2$, HRP oxidizes ABTS to a green-colored product exhibiting a characteristic absorbance maximum at 414 nm. Assay reactions were carried out in 96-well microtiter plates, and absorbance was recorded using a SPECTROstar Omega plate reader (BMG LABTECH). Each assay was performed in a total reaction volume of 250 µL. The standard reaction mixture contained 50 µM ABTS and 10 µg HRP (150–250 U $mg^{-1}$) in 100 mM $NaP_i$ buffer (pH 6.0). Prior to performing the ABTS–HRP assay, reactions were quenched by adding 1 M HCl at a 1:1 volumetric ratio to stop enzymatic activity.

### 2. Supporting Figures and Tables

#### 2.1 Supporting Tables

**Table S1.** GC-FID parameters used for analytics of indene (**1**) and products.

| **Gas chromatograph** | **Shimadzu GC-2010 Plus (Kyoto, Japan)** |
|---|---|
| **Column** | Optima™ 5 MS GC from Macherey-Nagel™ |
| **Length**<br>**Inner diameter**<br>**Film thickness** | 30 m<br>0.25 mm<br>0.25 µM |
| **Injection volume** | 1 µL |
| **Injection temp.** | 230 °C |
| **Injection mode** | Split |
| **Carrier gas** | $N_2$ |
| **Flow control mode** | Linear velocity |
| **Pressure** | 158.7 kPa |
| **Total flow** | 54.0 mL $min^{-1}$ |
| **Column flow** | 2.37 mL $min^{-1}$ |
| **Linear velocity** | 49.1 cm $s^{-1}$ |
| **Purge flow** | 3.0 mL $min^{-1}$ |
| **Split ratio** | 20.5 |
| **Oven temp. program** | 70 °C, 5 °C $min^{-1}$ to 110 °C, 15 °C $min^{-1}$ to 280 °C, hold for 5 min |
| **FID temperature** | 250 °C |
| **Retention times (RT)** | Indene (**1**): 5.82 min<br>acetophenone (IS): 6.19 min<br>1*H*-indenol (**1a**): 9.49 min<br>*cis*-indanediol (**1b**): 12.46 min |

**Table S2.** GC-FID parameters used for analytics of toluene (**2**) and products.

| **Gas chromatograph** | **Shimadzu GC-2010 Plus (Kyoto, Japan)** |
|---|---|
| **Column** | Optima™ 5 MS GC from Macherey-Nagel™ |
| **Length**<br>**Inner diameter**<br>**Film thickness** | 30 m<br>0.25 mm<br>0.25 µM |
| **Injection volume** | 1 µL |
| **Injection temp.** | 230 °C |
| **Injection mode** | Split |
| **Carrier gas** | $N_2$ |
| **Flow control mode** | Linear velocity |
| **Pressure** | 158.7 kPa |
| **Total flow** | 54.0 mL $min^{-1}$ |
| **Column flow** | 2.37 mL $min^{-1}$ |
| **Linear velocity** | 49.1 cm $s^{-1}$ |
| **Purge flow** | 3.0 mL $min^{-1}$ |

| | |
|---|---|
| **Split ratio** | 20.5 |
| **Oven temp. program** | 70 °C, 5 °C $min^{-1}$ to 90 °C, hold for 15 min, 20 °C $min^{-1}$ to 280 °C, hold for 5 min |
| **FID temperature** | 250 °C |
| **Retention times (RT)** | toluene (**2**): 2.13 min<br>benzyl alcohol (**2a**): 5.72 min<br>*o*-cresol (**2c**): 6.24 min<br>acetophenone (IS): 6.55 min<br>*m*-cresol (**2d**): 6.9 min<br>toluene dihydrodiol (**2b**): 7.15 min |

**Table S3.** GC-MS parameters used for toluene (**2**) and products detection.

| **Instrument** | **GCMS-QP2010 SE (Shimadzu Europa GmbH.)** |
|---|---|
| **Column** | Optima™ 5 MS GC from Macherey-Nagel™ |
| **Length**<br>**Inner diameter**<br>**Film thickness** | 30 m<br>0.25 mm<br>0.25 µM |
| **Injection volume** | 1 µL |
| **Injection temp.** | 250 °C |
| **Injection mode** | Split |
| **Carrier gas** | $N_2$ |
| **Flow control mode** | Linear velocity |
| **Pressure** | 116.3 kPa |
| **Total flow** | 51.9 mL $min^{-1}$ |
| **Column flow** | 1.88 mL $min^{-1}$ |
| **Linear velocity** | 50 cm $s^{-1}$ |
| **Purge flow** | 3.0 mL $min^{-1}$ |
| **Split ratio** | 25 |
| **Oven temp. program** | 60 °C, hold for 2 min, 5 °C $min^{-1}$ to 80 °C, hold for 15 min, 14 °C $min^{-1}$ to 300 °C, hold for 2 min |
| **MS parameters** | |
| **Ion source temp.** | 200 °C |
| **Interface temp.** | 250 °C |
| **Mode** | Scan |
| **Scan range** | 50–350 m/z |

**Table S4.** Oligonucleotides used in this study. Mutated triplet bases are highlighted in bold and olive green.

| **Name** | **Sequence 5'→ 3'** |
|---|---|
| **HMI_40_CDO_F_E25A-F** | aagcaaaggt**gca**gcagttgccatttttaac |
| **HMI_40_CDO_F_E25A-R** | tggcaactgc**tgc**acctttgctttcaacctg |
| **HMI_41_CDO_F_D41A-F** | ttgcaacccag**gca**cgttgtacccatggtg |
| **HMI_41_CDO_F_D41A-R** | gggtacaacg**tgc**ctgggttgcaaacagtt |
| **HMI_43_CDO_F_R42A-F** | aacccaggat**gca**tgtacccatggtgattgg |
| **HMI_43_CDO_F_R42A-R** | accatgggtaca**tgc**atcctgggttgcaaac |
| **HMI_42_CDO_F_D47A-F** | tacccatggt**gca**tggagcctgagcgaagg |
| **HMI_42_CDO_F_D47A-R** | aggctcca**tgc**accatgggtacaacgatc |
| **HMI_44_CDO_F_W48A-F** | acccatggtgat**gca**agcctgagcgaaggtgg |
| **HMI_44_CDO_F_W48A-R** | tcgctcaggct**tgc**atcaccatgggtacaacg |
| **HMI_39_CDO_F_E52A-F** | agcctgagc**gca**ggtggttatctggaag |
| **HMI_39_CDO_F_E52A-R** | ataaccacc**tgc**gctcaggctccaatcac |
| **HMI_185_CDO fd E62A-F** | gatattgtt**gca**tgtagcctgcatatgggtc |
| **HMI_185_CDO fd E62A-R** | aggctaca**tgc**aacaatatcaccttccagat |
| **HMI_67_CDO_E84A-F** | accgccttgt**gca**ccgctgaaaatctatccg |
| **HMI_67_CDO_E84A-R** | tttcagcgg**tgc**acaaggcggtgctgc |
| **HMI_66_CDO_K87A-F** | tgaaccgctg**gca**atctatccgattcgtattg |
| **HMI_66_CDO_K87A-R** | tcggatagat**tgc**cagcggttcacaaggcg |
| **HMI_146_NDO fd D42A-F** | tacgccacc**gca**aacctgtgcacgcatg |
| **HMI_146_NDO fd D42A-R** | tgcacaggttt**tgc**ggtggcgtagatttcccct |
| **HMI_166_NDO fd T46A-F** | aacctgtgc**gcg**catggttccgcgc |
| **HMI_166_NDO fd T46A-R** | aaccatg**cgc**gcacaggttgtcggtgg |
| **HMI_167_NDO fd H47A-F** | tgtgcacg**gcg**ggttccgcgc |
| **HMI_167_NDO fd H47A-R** | ggaacc**cgc**cgtgcacaggttgtcgg |
| **HMI_147_NDO fd D54A-F** | cgcatgtct**gca**ggctatctcgaggggcga |
| **HMI_147_NDO fd D54A-R** | tcgagatagcc**tgc**agacatgcgcgcggaa |
| **HMI_127_NDO fd R60A-F** | tctcgagggg**gcc**gaaatagaatgccc |
| **HMI_127_NDO fd R60A-R** | ttctatttc**ggc**cccctcgagatagccatc |
| **HMI_128_NDO fd E61A-F** | gaggggcga**gcc**atagaatgccctttg |
| **HMI_128_NDO fd E61A-R** | gcattctat**ggc**tcgcccctcgagatagc |
| **HMI_148_NDO fd E63A-F** | cgagaaata**gca**tgccctttgcatcaaggtcg |
| **HMI_148_NDO fd E63A-R** | caaagggca**tgc**tatttctcgcccctcg |
| **HMI_164_NDO fd L66A-F** | gaatgccct**gcc**catcaaggtcggtttgac |
| **HMI_164_NDO fd L66A-R** | accttgatg**ggc**agggcattctatttctcgcc |
| **HMI_149_NDO fd H67A-F** | tgccctttg**gca**caaggtcggtttgacg |
| **HMI_149_NDO fd H67A-R** | ccgaccttg**tgc**caaagggcattctatttctcgc |
| **HMI_129_NDO fd D72A-F** | ggtcggttt**gcc**gtttgcaccggtaaagcc |
| **HMI_129_NDO fd D72A-R** | ccggtgcaaac**ggc**aaaccgaccttgatgcaa |
| **HMI_165_NDO fd P82A-F** | ttatgcgca**gca**gtgacacagaacatcaaaac |
| **HMI_165_NDO fd P82A-R** | ctgtgtcac**tgc**tgcgcataaggctttac |
| **HMI_130_NDO fd K88A-F** | cagaacatc**gca**acatatcctgtgaagatcg |
| **HMI_130_NDO fd K88A-R** | aggatatgt**tgc**gatgttctgtgtcacagg |
| **HMI_126_NDO fd A50W-F** | atggttcc**tgg**cgcatgtctgatggc |
| **HMI_126_NDO fd A50W-R** | agacatgcg**cca**ggaaccatgcgtgc |

| | |
|---|---|
| **HMI_83_CDO_D41W-F** | tgcaacccag**tgg**cgttgtacccatgg |
| **HMI_83_CDO_D41W-R** | tacaacg**cca**ctgggttgcaaacagttcac |
| **HMI_357_CDOfd_D41R_F** | ttgcaacccag**cgt**cgttgtacccatggtg |
| **HMI_357_CDOfd_D41R_R** | gggtacaacg**acg**ctgggttgcaaacagtt |
| **HMI_358_CDOfd_D41N_F** | ttgcaacccag**aac**cgttgtacccatggtg |
| **HMI_358_CDOfd_D41N_R** | gggtacaacg**gtt**ctgggttgcaaacagtt |
| **HMI_359_CDOfd_W48F_F** | acccatggtgat**ttt**agcctgagcgaaggtgg |
| **HMI_359_CDOfd_W48F_R** | tcgctcaggct**aaa**atcaccatgggtacaacg |
| **HMI_360_CDOfd_W48E_F** | acccatggtgat**gaa**agcctgagcgaaggtgg |
| **HMI_360_CDOfd_W48E_R** | tcgctcaggct**ttc**atcaccatgggtacaacg |
| **HMI_361_CDOfd_W48H_F** | acccatggtgat**cat**agcctgagcgaaggtgg |
| **HMI_361_CDOfd_W48H_R** | tcgctcaggct**atg**atcaccatgggtacaacg |
| **HMI_362_CDOfd_W48S_F** | acccatggtgat**agc**agcctgagcgaaggtgg |
| **HMI_362_CDOfd_W48S_R** | tcgctcaggct**gct**atcaccatgggtacaacg |
| **HMI_381_CDOfd_E52K_F** | agcctgagc**aaa**ggtggttatctggaag |
| **HMI_381_CDOfd_E52K_R** | ataaccacc**ttt**gctcaggctccaatcac |
| **HMI_382_CDOfd_E62K_F** | gatattgtt**aaa**tgtagcctgcatatgggtc |
| **HMI_382_CDOfd_E62K_R** | aggctaca**ttt**aacaatatcaccttccagat |
| **HMI_99_CDO_K87E-F** | gaaccgctg**gaa**atctatccgattcgtattg |
| **HMI_99_CDO_K87E-R** | tcggatagat**ttc**cagcggttcacaaggcgg |
| **HMI_363_CDOfd_K87Q_F** | tgaaccgctg**cag**atctatccgattcgtattg |
| **HMI_363_CDOfd_K87Q_R** | tcggatagat**ctg**cagcggttcacaaggcg |
| **HMI_374_CDOfd_E84L_F** | accgccttgt**ctg**ccgctgaaaatctatccg |
| **HMI_374_CDOfd_E84L_R** | tttcagcgg**cag**acaaggcggtgctgc |
| **HMI_375_CDOfd_E84F_F** | accgccttgt**ttt**ccgctgaaaatctatccg |
| **HMI_375_CDOfd_E84F_R** | tttcagcgg**aaa**acaaggcggtgctgc |
| **HMI_376_CDOfd_E84Q_F** | accgccttgt**cag**ccgctgaaaatctatccg |
| **HMI_376_CDOfd_E84Q_R** | tttcagcgg**ctg**acaaggcggtgctgc |
| **HMI_100_CDO_E84K-F** | accgccttgt**aaa**ccgctgaaaatctatccg |
| **HMI_100_CDO_E84K-R** | ttttcagcgg**ttt**acaaggcggtgctgctttaac |
| **HMI_101_CDO_E84Y-F** | accgccttgt**tat**ccgctgaaaatctatcc |
| **HMI_101_CDO_E84Y-R** | tttcagcgg**ata**acaaggcggtgctgctttaac |
| **HMI_364_NDOfd_A50V_F** | atggttcc**gtg**cgcatgtctgatggc |
| **HMI_364_NDOfd_A50V_R** | agacatgcg**cac**ggaaccatgcgtgc |
| **HMI_365_NDOfd_A50L_F** | atggttcc**ctt**cgcatgtctgatggc |
| **HMI_365_NDOfd_A50L_R** | agacatgcg**aag**ggaaccatgcgtgc |
| **HMI_366_NDOfd_A50E_F** | atggttcc**gag**cgcatgtctgatggc |
| **HMI_366_NDOfd_A50E_R** | agacatgcg**ctc**ggaaccatgcgtgc |
| **HMI_367_NDOfd_A50H_F** | atggttcc**cat**cgcatgtctgatggc |
| **HMI_367_NDOfd_A50H_R** | agacatgcg**atg**ggaaccatgcgtgc |
| **HMI_368_NDOfd_A50S_F** | atggttcc**tct**cgcatgtctgatggc |
| **HMI_368_NDOfd_A50S_R** | agacatgcg**aga**ggaaccatgcgtgc |
| **HMI_380_NDOfd_D54K_F** | cgcatgtct**aag**ggctatctcgaggggcga |
| **HMI_380_NDOfd_D54K_R** | tcgagatagcc**ctt**agacatgcgcgcggaa |
| **HMI_369_NDOfd_E63L_F** | cgagaaata**ttg**tgccctttgcatcaaggtcg |
| **HMI_369_NDOfd_E63L_R** | caaagggca**caa**tatttctcgcccctcg |
| **HMI_370_NDOfd_E63F_F** | cgagaaata**ttt**tgccctttgcatcaaggtcg |
| **HMI_370_NDOfd_E63F_R** | caaagggca**aaa**tatttctcgcccctcg |

| | |
|---|---|
| **HMI_371_NDOfd_E63Q_F** | cgagaaatacagtgccctttgcatcaaggtcg |
| **HMI_371_NDOfd_E63Q_R** | caaagggcactgtatttctcgcccctcg |
| **HMI_372_NDOfd_E63K_F** | cgagaaataaagtgccctttgcatcaaggtcg |
| **HMI_372_NDOfd_E63K_R** | caaagggcactttatttctcgcccctcg |
| **HMI_373_NDOfd_E63Y_F** | cgagaaatatactgccctttgcatcaaggtcg |
| **HMI_373_NDOfd_E63Y_R** | caaagggcagtatatttctcgcccctcg |
| **HMI_299_CDO_E32A_F** | gtggatgaagccaaaggtctgttagatccgc |
| **HMI_299_CDO_E32A_R** | cagacctttggcttcatccaccagggc |
| **HMI_322_CDO_F144Y_F** | aaagaagcctattgcgataaaaaagaaggtg |
| **HMI_322_CDO_F144Y_R** | tttatcgcaataggcttctttttcatacgg |
| **HMI_295_CDO_K147A_F** | ttctgcgatgccaaagaaggtgattgcgg |
| **HMI_295_CDO_K147A_R** | accttctttggcatcgcagaaggcttcttt |
| **HMI_321_CDO_F154A_F** | gattgcggtgcagataaagcagattggg |
| **HMI_321_CDO_F154A_R** | tgctttatctgcaccgcaatcaccttctttt |
| **HMI_319_CDO_W159A_F** | aaagcagatgcaggtccgctgcaggc |
| **HMI_319_CDO_W159A_R** | agcggacctgcatctgctttatcgaaaccg |
| **HMI_294_CDO_D158A_F** | gataaagcagcatggggtccgctgcagg |
| **HMI_294_CDO_D158A_R** | cggaccccatgctgctttatcgaaaccgc |
| **HMI_231_CDO_R407A_F** | gcacgtagcgcaccgctgtgtgcacag |
| **HMI_231_CDO_R407A_R** | acacagcggtgcgctacgtgctttataacc |
| **HMI_232_CDO_E436A_F** | tatagcgaagcagcagcacgcggttttt |
| **HMI_232_CDO_E436A_R** | gcgtgctgctgcttcgctataaacatagctc |
| **HMI_323_CDO_A415F_F** | cagatgggtttcggtgttccgaataaaaacaa |
| **HMI_323_CDO_A415F_R** | ggaacaccgaaacccatctgtgcacaca |
| **HMI_211_CDO oxy R110A-F** | cgtattgaagccagcgattttggtaatgcc |
| **HMI_211_CDO oxy R110A-R** | aaaatcgctggcttcaatacgcataccacg |
| **HMI_3_CDO_A-K117A-F** | tggtaatgccgcaagctttacctgtaccta |
| **HMI_3_CDO_A-K117A-R** | aggtaaagcttgcggcattaccaaaatcgc |
| **HMI-24_CDO_D129A-F** | tgggcatatgcaaccgcaggtaatctgg |
| **HMI-24_CDO_D129A-R** | acctgcggttgcatatgcccaaccatgatag |
| **HMI_7_CDO_B-R65A-F** | acataccagcgcaaataaagccatggaatatg |
| **HMI_7_CDO_B-R65A-R** | ggctttatttgcgctggtatgtgtggtacg |
| **HMI_8_CDO_B-W100A-F** | ggtctgaatgccaccgaagatccgcct |
| **HMI_8_CDO_B-W100A-R** | atcttcggtggcattcagaccgctaacac |
| **HMI_233_CDO_E102A_F** | aattggaccgccgatccgcctagccgt |
| **HMI_233_CDO_E102A_R** | aggcggatcggcggtccaattcagacc |
| **HMI_234_CDO_D103A_F** | tggaccgaagcaccgcctagccgtag |
| **HMI_234_CDO_D103A_R** | gctaggcggtgcttcggtccaattcagaccg |
| **HMI_383_TDOxy V99K-F** | taccagcgataaatcctggagcgaaaatcc |
| **HMI_383_TDOxy V99K-R** | gctccaggatttatcgctggtaattttgcg |
| **HMI_113_CDOfd vector-F** | accggtaaagttaaagcagcaccgcc |
| **HMI_113_CDOfd vector-R** | attctgggttgcaaacagttcaccatcaacg |
| **HMI_121_NDO motif-F** | gaactgtttgcaacccagaacctgtgcacgcatggttcc |
| **HMI_121_NDO motif-R** | gctgctttaactttaccggtgcaaacgtcaaaccgaccttgatgc |
| **HMI_122_CDO motif-F** | aatctacgccaccgacgatcgttgtacccatggtgattggag |
| **HMI_122_CDO motif-R** | gcataaggctttaccggtacgaacacaaaaacgacccatatgcag |
| **HMI_123_NDOfd vector-F** | accggtaaagccttatgcgcacctgt |
| **HMI_123_NDOfd vector-R** | gtcggtggcgtagatttccccttccac |

| | |
|---|---|
| **HMI_124_NDO fd S49D-F** | acgcatggtgatgcgcgcatgtctgatg |
| **HMI_124_NDO fd S49D-R** | catgcgcgcatcaccatgcgtgcacagg |
| **HMI_125_NDO fd P65S Q68M-F** | aaatagaatgcagtttgcatatgggtcggtttgacgtttgcacc |
| **HMI_125_NDO fd P65S Q68M-R** | aaaccgacccatatgcaaactgcattctatttctcgcccctcgag |
| **HMI_116_CDO fd D47S-F** | acccatggtagctggagcctgagcgaagg |
| **HMI_116_CDO fd D47S-R** | caggctccagctaccatgggtacaacgatcc |
| **HMI_117_CDO fd S64P M67Q-F** | gttgaatgtccgctgcatcagggtcgtttttgtgttcgtacc |
| **HMI_117_CDO fd S64P M67Q-R** | aaaaacgaccctgatgcagcggacattcaacaatatcaccttccag |
| **HMI_300_bnzAB_F** | atgggtcgcggatccatgaaccagaccgatacctcgccg |
| **HMI_300_bnzAB_R** | ggtggtggtgctcgagtcaaaagaaaaagctcaggttattggccagaatgg |
| **HMI_310_todA_F** | tgggtcgcggatccatggcaactcacgtagcaataataggtaat |
| **HMI_310_todA_R** | ggtggtggtgctcgagttacgtgaggtccccttcgttt |
| **HMI_311_todB_F** | tgggtcgcggatccatgacctggacctatattctgcgc |
| **HMI_311_todB_R** | ggtggtggtgctcgagtcatttcagttcgccattatccagatcc |
| **HMI_312_vec_F** | ctcgagcaccaccaccaccacc |
| **HMI_312_vec_R** | ggatccgcgacccatttgctgtcc |

**Table S5: Determined ECNs and RFs of indanone and 1*H*-indenol.**

| **Compound** | **ECN** | **RF** |
|---|---|---|
| indanone | 8.20[a] | 1.00 |
| 1*H*-indenol | 8.42[a] | 1.03 |

[a] ECNs were calculated based on the model of Jorgensen et al[5].

**Table S6.** Rationally designed CDO-Oxy variants generated in this study.

| CDO-Oxy variant | Location | Subunit | Purpose |
|---|---|---|---|
| **E32A** | Top surface | α-Subunit | Minimize electrostatic interaction |
| **F144Y** | Top surface | α-Subunit | Emulate NDO-Oxy |
| **K147A** | Top surface | α-Subunit | Minimize electrostatic interaction |
| **F154A** | Top surface | α-Subunit | Minimize hydrophobic interaction |
| **D158A** | Top surface | α-Subunit | Minimize electrostatic interaction |
| **W159A** | Top surface | α-Subunit | Minimize hydrophobic interaction |
| **R407A** | Top surface | α-Subunit | Minimize electrostatic interaction |
| **E436A** | Top surface | α-Subunit | Minimize electrostatic interaction |
| **A415F** | Top surface | α-Subunit | Emulate NDO-Oxy |
| **R110A** | Side surface | α-Subunit | Minimize electrostatic interaction |
| **K117A** | Side surface | α-Subunit | Minimize electrostatic interaction |
| **D129A** | Side surface | α-Subunit | Minimize electrostatic interaction |
| **R65A** | Side surface | β-Subunit | Minimize electrostatic interaction |
| **L98K** | Side surface | β-Subunit | Unintentionally generated during mutagenesis |
| **W100A** | Side surface | β-Subunit | Minimize hydrophobic interaction |
| **E102A** | Side surface | β-Subunit | Minimize electrostatic interaction |
| **D103A** | Side surface | β-Subunit | Minimize electrostatic interaction |

**Table S7.** EPR parameters Rieske clusters of the different Fd and Oxy variants.

| | $g_z$ | $g_y$ | $g_x$ |
|---|---|---|---|
| **NDO Fd** | 2.020 | 1.906 | 1.813 |
| **NDO Fd A50W** | 2.015 | 1.911 | 1.801 |
| **CDO Fd** | 2.021 | 1.902 | 1.823 |
| **CDO Oxy** | 2.023 | 1.911 | 1.760 |
| **CDO Oxy L98K** | 2.023 | 1.911 | 1.760 |

**Table S8.** Statistical analysis of Fd-Oxy complex docking result acquired by HADDOCK.

| Analysis component | NDO-Fd A50W/CDO-Oxy | NDO-Fd A50W/CDO-Oxy β_L98K |
|---|---|---|
| **HADDOCK score** | -60.8 +/- 8.6 | -73.0 +/- 6.0 |
| **RMSD from the overall lowest-energy structure** | 3.5 +/- 0.2 | 4.7 +/- 0.1 |
| **Van der Waals energy (kcal mol$^{-1}$)** | -36.8 +/- 9.0 | -30.0 +/- 5.0 |
| **Electrostatic energy (kcal mol$^{-1}$)** | -115.5 +/- 16.2 | -228.2 +/- 14.3 |
| **Desolvation energy (kcal mol$^{-1}$)** | -10.5 +/- 1.3 | 1.8 +/- 2.1 |
| **Restraints violation energy (kcal mol$^{-1}$)** | 95.5 +/- 1.8 | 8.3 +/- 1.6 |
| **Buried surface area (Å$^2$)** | 1397.0 +/- 129.6 | 1272.0 +/- 46.8 |
| **Z-score** | -1.1 | -1.4 |

#### 2.2 Supporting Schemes and Figures

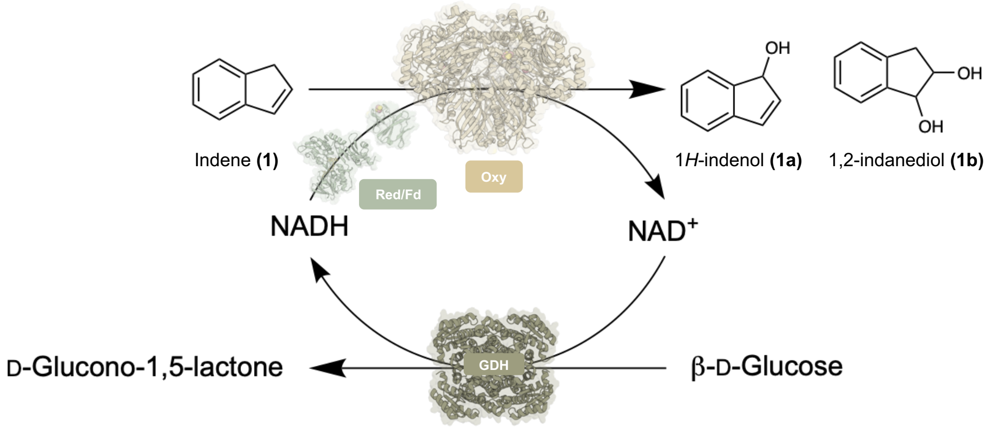

**Scheme S1.** *In vitro* conversion of indene (**1**) to 1*H*-indenol (**1a**) and 1,2-indandiol (**1b**) catalyzed by the RO. Glucose dehydrogenase (GDH) is used for the regeneration of the NADH cofactor.

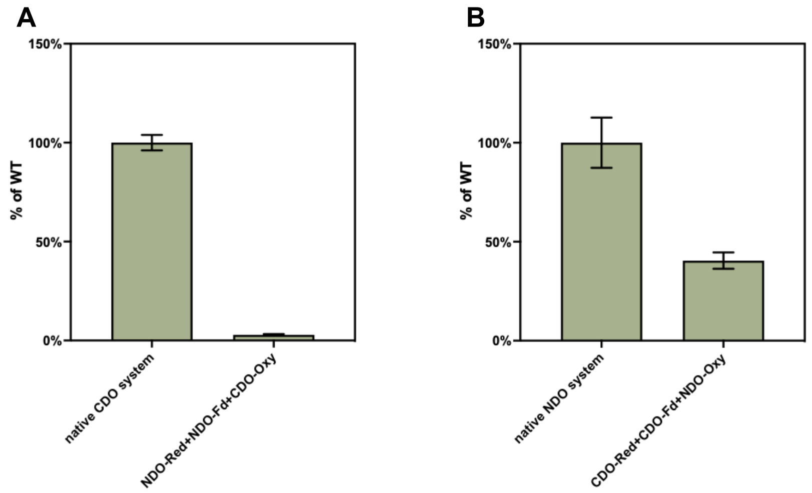

**Figure S1.** Assessment of the cross-reactivity of (A) NDO-Oxy and (B) CDO-Oxy. The product concentrations of native systems were normalized to 100%. Reaction conditions: 4 µM Oxy, 30 µM Fd, 10 µM Red, 10 mM **1**, 50 mM D-glucose, 10 U GDH, 400 µM $NAD^+$, 1 mg $mL^{-1}$ catalase, 1 mM DTT, 50 mM $NaP_i$ (pH 7.2), 30 °C, 120 rpm, 3 h. Product formation was analyzed by GC-FID. Product concentrations correspond to the combined amounts of **1a** and **1b**. The data points represent the mean, and the error bars represent the standard deviation of duplicate experiments (n = 2).

```
CARDO-FdIIB   MNRHSAGQSTPVRVATLDQLKPGVPTAFDVDG-DEVMVVRDGDSVYAISNLCSHAEAYLD 59
CARDO-FdIII   ------MNQIWLKVCAASDMQPGTIRRVNRVGAAPLAVYRVGDQFYATEDTCTHGIASLS 54
CDO-Fd        --------MTFSKVCEVSDVPVGDALQVESKG-EAVAIFNVDGELFATQDRCTHGDWSLS 51
NDO-Fd        ------MTVKWIEAVALSDILEGDVLGVTVEG-KELALYEVEGEIYATDNLCTHGSARMS 53
                            ..   .::  *     .    *    : : .   ...:* .: *:*.    :.

CARDO-FdIIB   MGVFHAESLEIECPLHVGRFDVRTGAPTALPCVLPVRAYDVVVDGTEILVAPKEAD---  115
CARDO-FdIII   -EGT-LDGDVIECPFHGGAFNVCTGMPASSPCTVPLGVFEVEVKEGEVYVAGEKK----  107
CDO-Fd        EGGY-LEGDIVECSLHMGRFCVRTGKVKAAPPCEPLKIYPIRIDGSDVFVDFDAGYLAP  109
NDO-Fd        -DGY-LEGREIECPLHQGRFDVCTGKALCAPVTQNIKTYPVKIENLRVMIDLS------  104
                    :.   :** :* * * * **    . *     :   : : :.    : :   .
```

**Figure S2.** Amino acid sequence alignments of CARDO-Fd$_{IIB}$ (UniProt ID: Q2HWH6), CARDO-Fd$_{III}$ (UniProt ID: Q8GI16), CDO-Fd (UniProt ID: Q51746) and NDO-Fd (UniProt ID: P0A185). Corresponding residues previously identified in CARDO are highlighted in yellow, while additional surface residues are shown in pink. Multiple sequence alignments were performed using Clustal Omega[6]. Default settings were kept for all parameters.

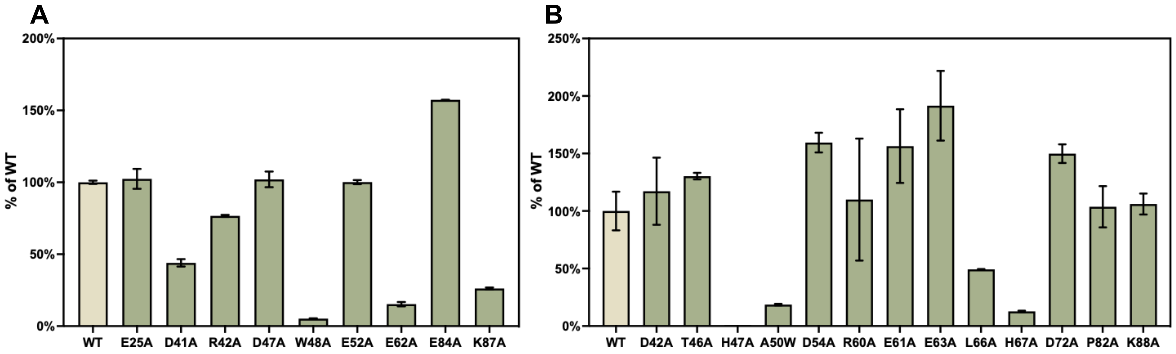

**Figure S3.** Catalytic activities of alanine-scanning variants of (A) CDO-Fd and (B) NDO-Fd. Wild-type (WT, product concentration normalized to 100%) is displayed as a beige bar, with green bars indicating the relative product concentrations of each variant. Reaction conditions: 4 µM Oxy, 30 µM Fd, 10 µM Red, 10 mM **1**, 50 mM D-glucose, 10 U GDH, 400 µM $NAD^+$, 1 mg $mL^{-1}$ catalase, 1 mM DTT, 50 mM $NaP_i$ (pH 7.2), 30 °C, 120 rpm, 3 h. Product formation was analyzed by GC-FID. Product concentrations correspond to the combined amounts of **1a** and **1b**. The data points represent the mean, and the error bars represent the standard deviation of duplicate experiments (n = 2).

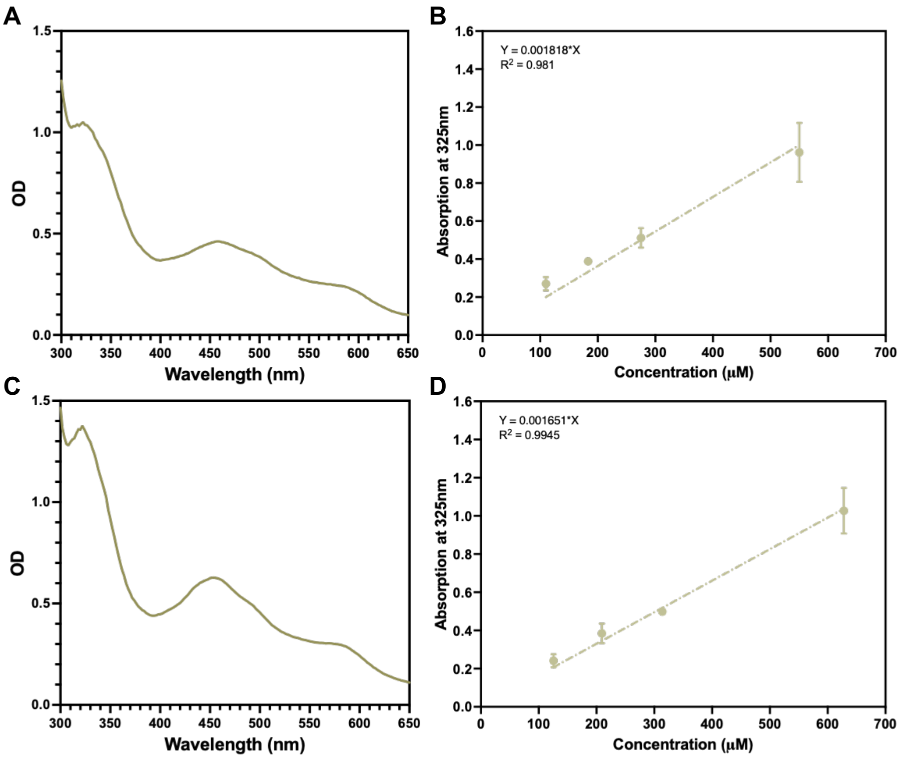

**Figure S4.** UV/Vis spectroscopic characterization and quantitative calibration of Fd chimeras and NDO-Fd A50W. (A) UV/VIS spectrum of purified CDO-Fd within a range of 300–650 nm. The measurement was performed using 550 μM of purified CDO-Fd in 50 mM $NaP_i$ (pH 7.2). (B) Standard curve for determining the concentration of Fds. Formula: y = 0.001818*x. Mean ± SD; n = 3. (C) UV/VIS spectrum of purified NDO-Fd within a range of 300–650 nm. The measurement was performed using 628 μM of purified NDO-Fd in 50 mM $NaP_i$ (pH 7.2). (D) Standard curve for determining the concentration of Fds. Formula: y = 0.001651*x. Mean ± SD; n = 3.

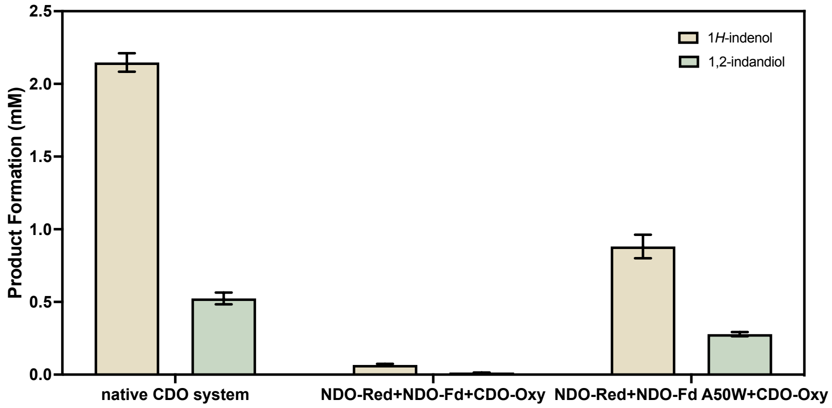

**Figure S5.** Assessment of NDO–CDO hybrid system activity. Product **1a** is shown in beige, and **1b** is shown in green. Reaction conditions: 4 μM Oxy, 30 μM Fd, 10 μM Red, 10 mM **1**, 50 mM D-glucose, 10 U GDH, 400 μM $NAD^+$, 1 mg $mL^{-1}$ catalase, 1 mM DTT, 50 mM $NaP_i$ (pH 7.2), 30 °C, 120 rpm, 3h. Product formation was analyzed by GC-FID. The data points represent the mean, and the error bars represent the standard deviation of duplicate experiments (n = 2).

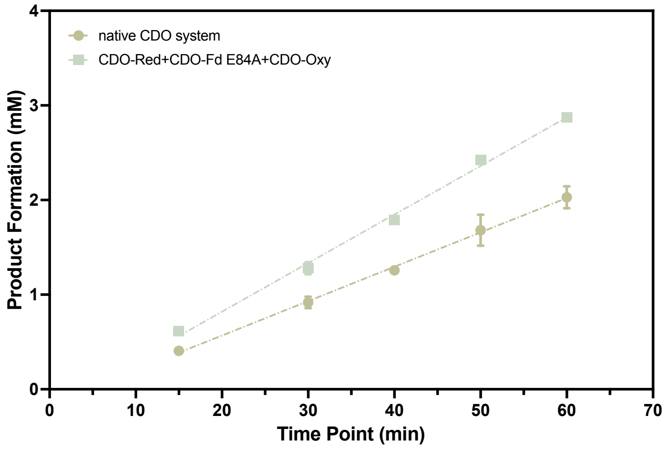

**Figure S6.** Time course analysis of total product formation in reactions catalyzed by the native CDO system or CDO-Red + CDO-Fd E84A + CDO-Oxy. Reaction conditions: 4 µM Oxy, 30 µM Fd, 10 µM Red, 10 mM **1**, 50 mM D-glucose, 10 U GDH, 400 µM $NAD^+$, 1 mg $mL^{-1}$ catalase, 1 mM DTT, 50 mM $NaP_i$ (pH 7.2), 30 °C, 120 rpm. Product formation was analyzed by GC-FID. Product concentrations correspond to the combined amounts of **1a** and **1b**. The data points represent the mean, and the error bars represent the standard deviation of duplicate experiments (n = 2).

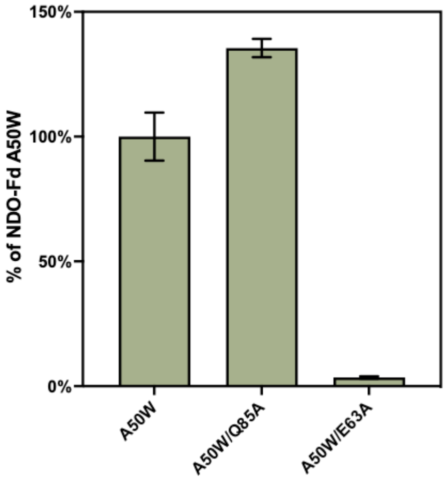

**Figure S7.** Activity assessment of the NDO–CDO hybrid system with different NDO-Fd variants. The product concentration of variant A50W was normalized to 100%, and other variants show relative product concentrations of variants A50W. Reaction conditions: 4 µM Oxy, 30 µM Fd, 10 µM Red, 10 mM **1**, 50 mM D-glucose, 10 U GDH, 400 µM $NAD^+$, 1 mg $mL^{-1}$ catalase, 1 mM DTT, 50 mM $NaP_i$ (pH 7.2), 30 °C, 120 rpm, 3h. Product formation was analyzed by GC-FID. Product concentrations correspond to the combined amounts of **1a** and **1b**. The data points represent the mean, and the error bars represent the standard deviation of duplicate experiments (n = 2).

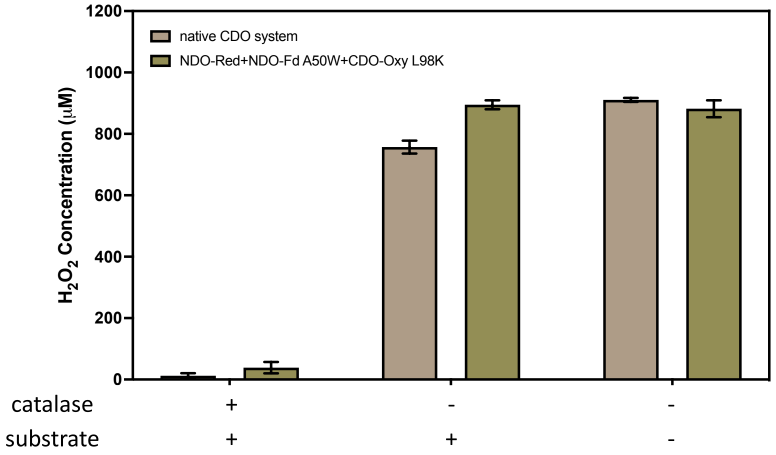

**Figure S8.** The amount of $H_2O_2$ generated after 2h in reactions catalyzed by the native CDO system or the NDO-Red + NDO-Fd A50W + CDO-Oxy L98K hybrid system. The data points represent the mean, and the error bars represent the standard deviation of duplicate experiments (n = 2).

**A**

```
TDO-Fd      MTWTYILRQGDLPPGEMQRYEGGPEPVMVCNVDGEFFAVQDTCTHGDWALSDG-YLDGDI 59
CDO-Fd      MTFSKVCEVSDVPVGDALQVESKGEAVAIFNVDGELFATQDRCTHGDWSLSEGGYLEGDI 60
            **:: : . .*:* *:  : *.  * * : *****:**.** ******:**:* **:***

TDO-Fd      VECTLHFGKFCVRTGKVKALPACKPIKVFPIKVEGDEVHVDLDNGELK- 107
CDO-Fd      VECSLHMGRFCVRTGKVKAAPPCEPLKIYPIRIDGSDVFVDFDAGYLAP 109
            ***:**:*:********** * *:*:*::**:::*.:*.**:* * *
```

**B**

```
TDO-Oxy     MIDSANRADVFLRKPAPVAPELQHEVEQFYYWEAKLLNDRRFEEWFALLAEDIHYFMPIR 60
CDO-Oxy     -MTSADLTKPIEWPEMPVSLELQNAVEQFYYREAQLLDYQNYEAWLALLTQDIQYWMPIR 59
             : **: :. :     **: ***: ****** **:**: :.:* *:***::**:*:****

TDO-Oxy     TTRIMRDSRLEYSGSREYAHFDDDATMMKGRLRKITSDVSWSENPASRTRHLVSNVMIVG 120
CDO-Oxy     TTHTSRNKAMEYVPPGGNAHFDETYESMRARIRARVSGLNWTEDPPSRSRHIVSNVIVRE 119
            **:  *:. :**      ****:    *:.*:*  .*.:.*:*:* **:**:****::

TDO-Oxy     AEAEGEYEISSAFIVYRNRLERQLDIFAGERRDTLRRNTSEAGFEIVNRTILIDQSTILA 180
CDO-Oxy     TESAGTLEVSSAFLCYRNRLERMTDIYVGERRDILLRVSDGLGFKIAKRTILLDQSTITA 179
            :*: *  *:****: *******  **:.***** * * :.  **:*.:****:***** *

TDO-Oxy     NNLSFFF 187
CDO-Oxy     NNLSQFF 186
            **** **
```

**Figure S9.** Amino acid sequence alignments of (A) TDO-Fd and CDO-Fd, and (B) β-subunit of TDO-Oxy and CDO-Oxy. Critical residues are marked in green. Multiple sequence alignments were performed using Clustal Omega[6]. Default settings were kept for all parameters.

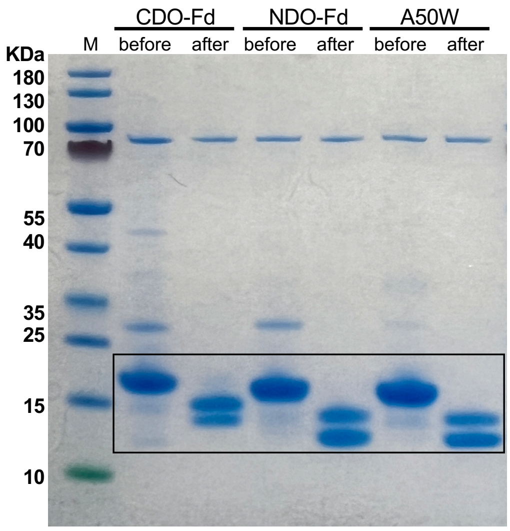

**Figure S10.** SDS–PAGE of Fds before and after N-terminal His-tag. CDO-Fd shows a molecular weight of 15.333 kDa before cleavage, and 13.451 kDa and 11.933 kDa after cleavage. NDO-Fd shows a molecular weight of 14.990 kDa before cleavage, and 13.107 kDa and 11.590 kDa after cleavage. The two bands after cleavage arise from two N-terminal thrombin recognition sites. M: PageRuler$^{TM}$ prestained protein ladder (Invitrogen by Thermo Scientific).

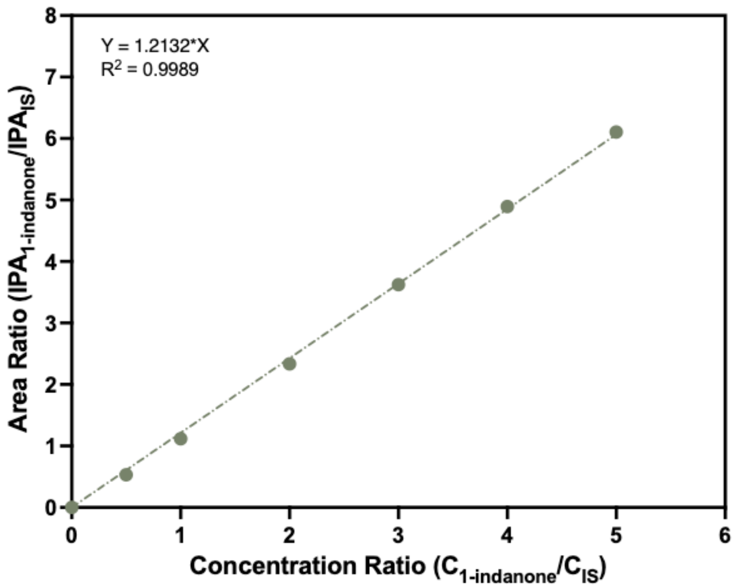

**Figure S11.** Standard curve for 1-indanone using acetophenone as internal standard (IS) analyzed by non-chiral GC-FID. The linear regression is shown. IPA denotes integrated peak area.

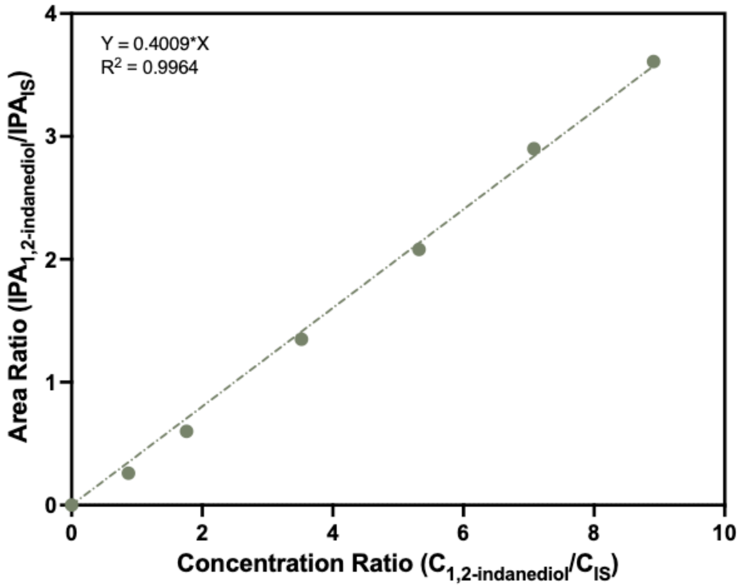

**Figure S12.** Standard curve for *cis*-1,2-indanediol using acetophenone as internal standard (IS) analyzed by non-chiral GC-FID. The linear regression is shown. IPA denotes integrated peak area.

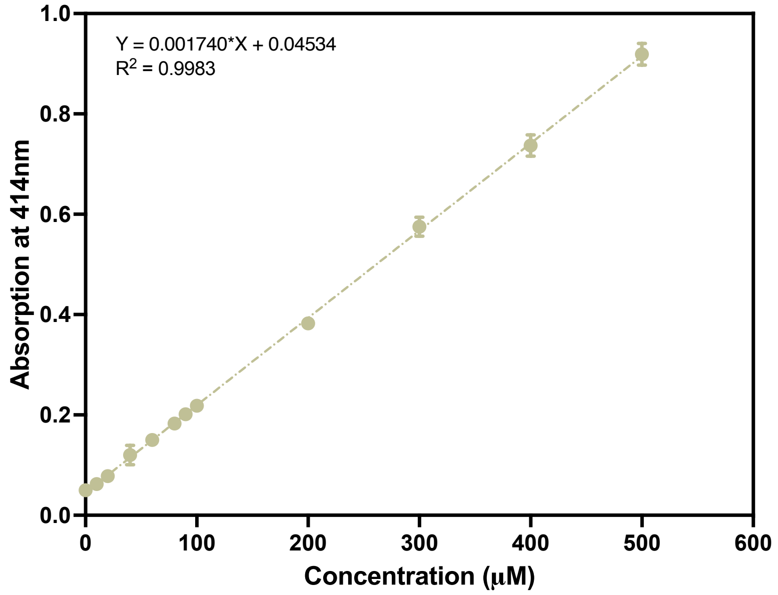

**Figure S13.** Standard curve for $H_2O_2$ determination via ABTS-HRP assay measuring the absorption at 414 nm. The linear regression is shown. The data points represent the mean, and the error bars represent the standard deviation of triplicate experiments (n = 3).

#### 3. Example chromatograms from non-chiral GC-FID analysis

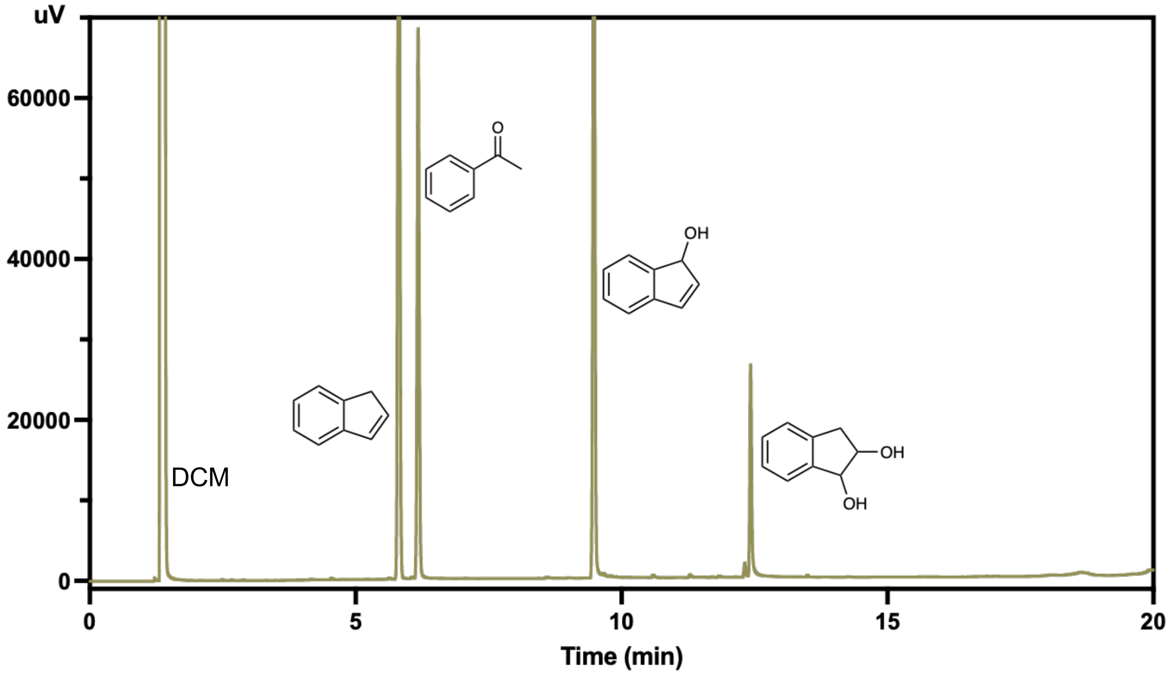

**Figure S14.** GC–FID chromatogram of a representative reaction mixture containing solvent DCM, internal standard, indene and related products. RT = 5.82 min for indene; 6.19 min for acetophenone (IS); 9.49 min for 1*H*-indenol; 12.46 min for *cis*-indanediol.

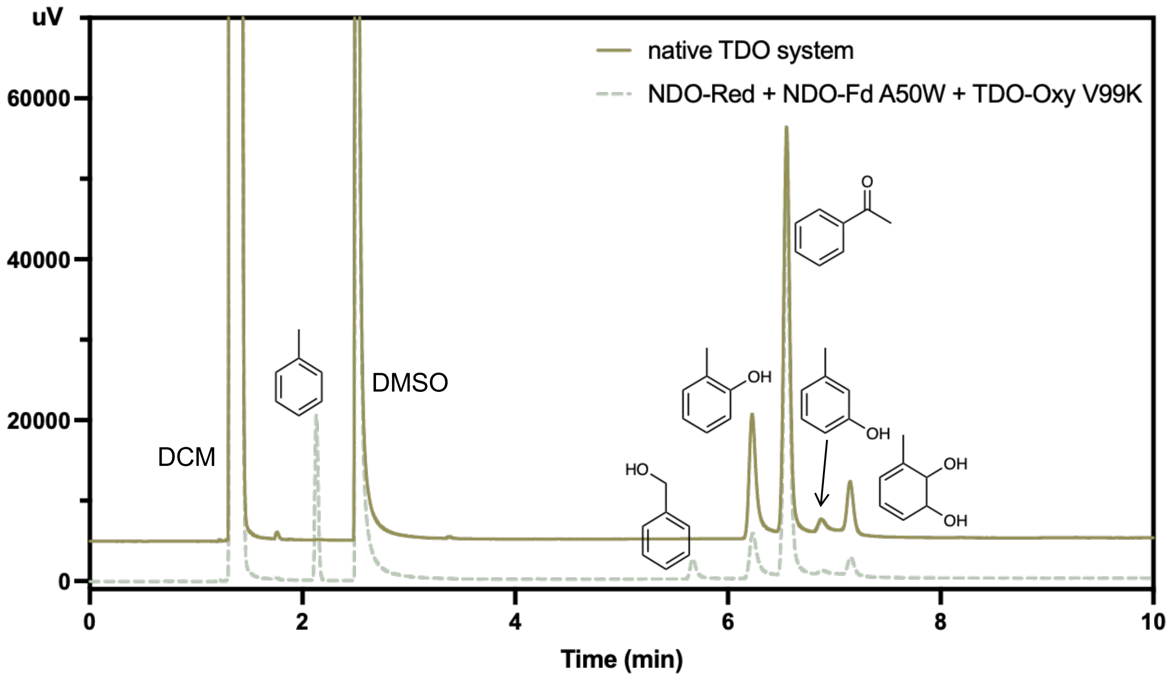

**Figure S15.** Stacked GC-FID chromatograms of reactions catalyzed by the native TDO system and the hybrid NDO-Red + NDO-Fd A50W + TDO-Oxy V99K system. RT = 2.13 min for toluene; 5.72 min for benzyl alcohol; 6.24 min for *o*-cresol; 6.55 min for acetophenone (IS); 6.9 min for *m*-cresol; 7.15 min for toluene dihydrodiol. DCM: dichloromethane.

#### 4. GC-MS chromatograms

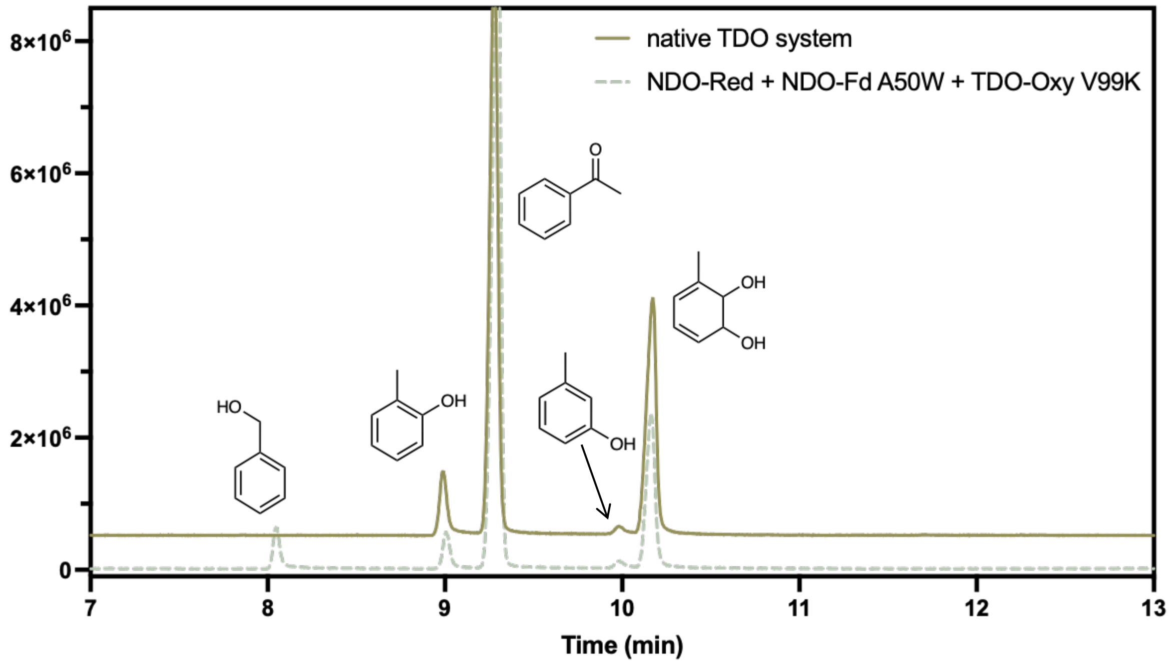

**Figure S16.** Stacked GC-MS chromatograms of reactions catalyzed by the native TDO system and the hybrid NDO-Red + NDO-Fd A50W + TDO-Oxy V99K system.

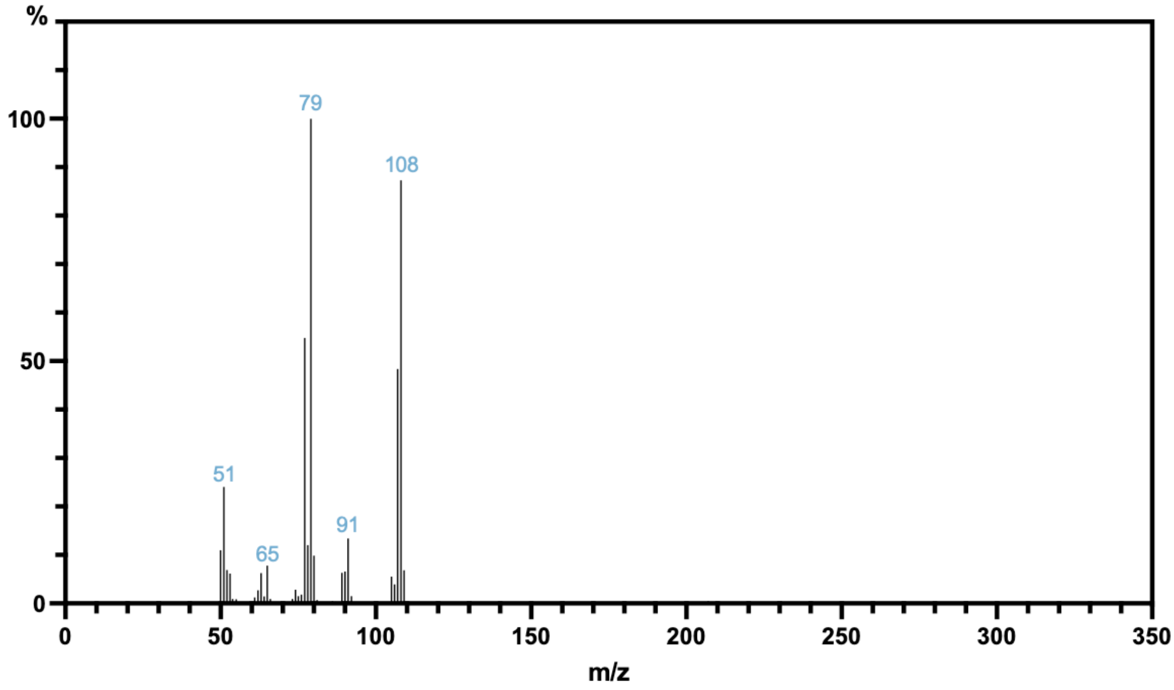

**Figure S17.** Mass spectrum for the peak at 8.05 min, consistent with reported data for benzyl alcohol[7].

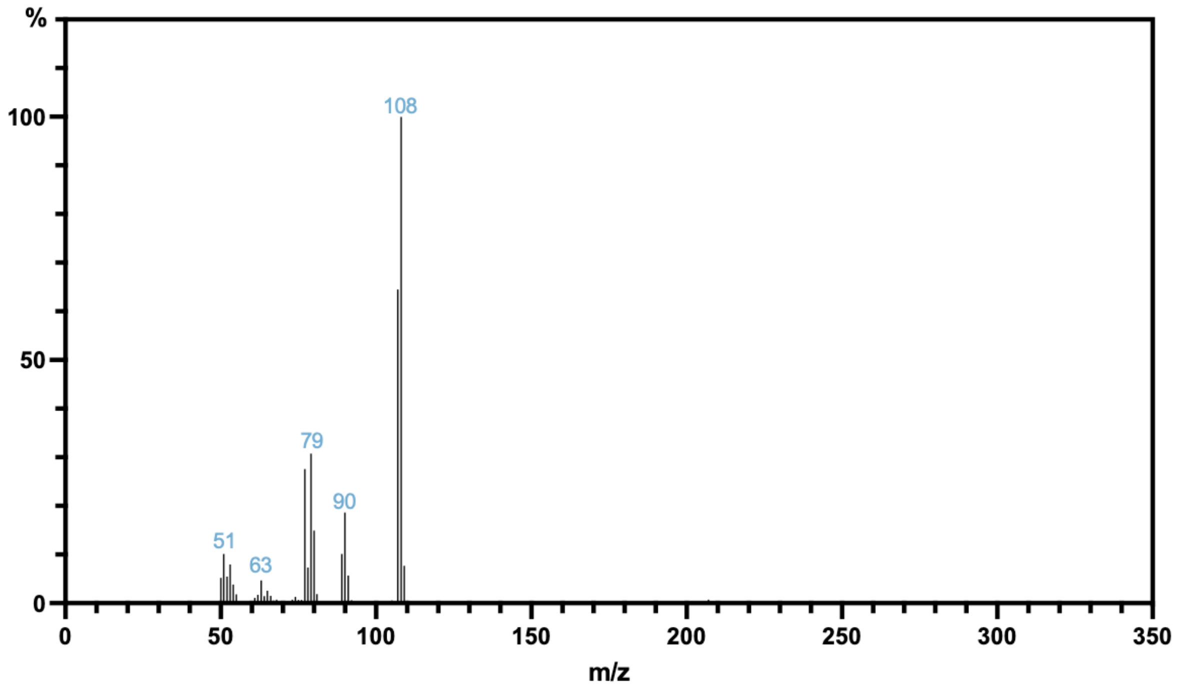

**Figure S18.** Mass spectrum for the peak at 9 min, consistent with reported data for *o*-cresol[7].

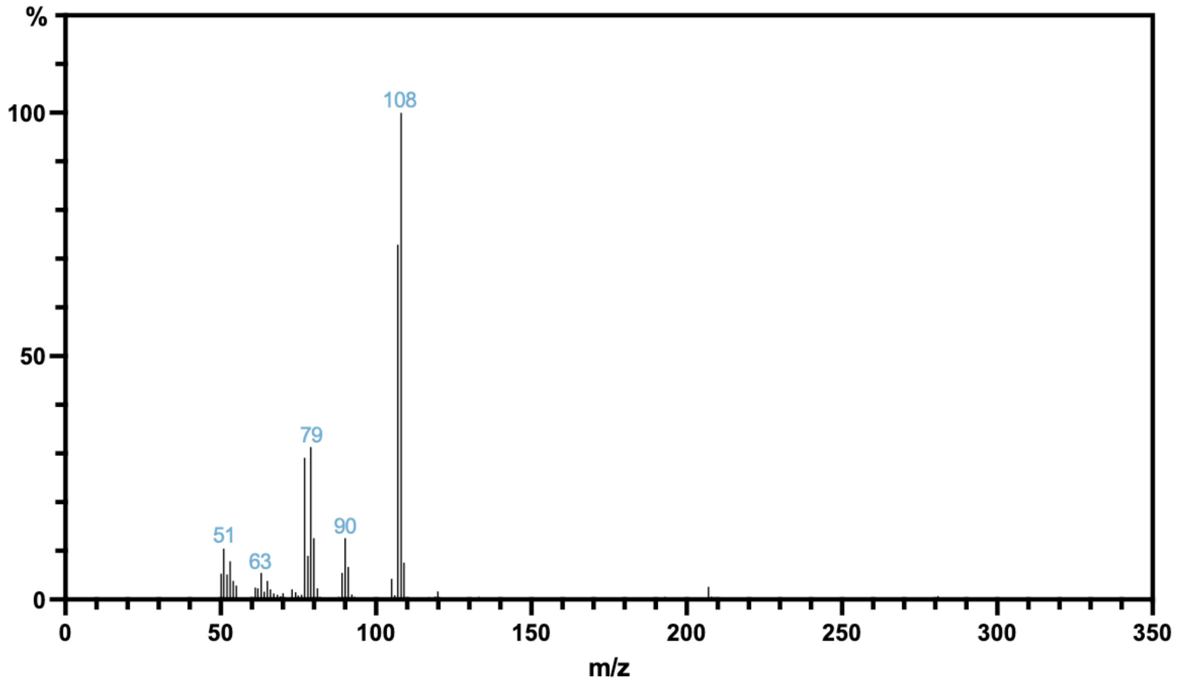

**Figure S19.** Mass spectrum for the peak at 9.97 min, consistent with reported data for *m*-cresol[7].

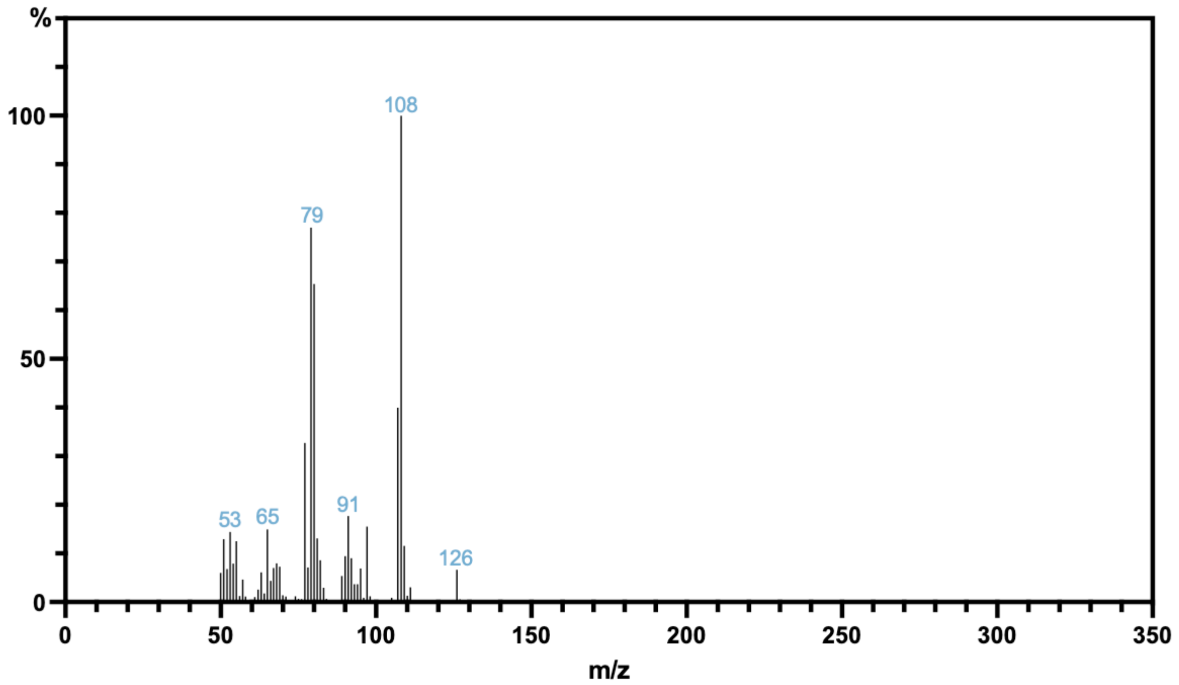

**Figure S20.** Mass spectrum for the peak at 10.15 min, consistent with reported data for toluene dihydrodiol[7].

#### 5. EPR spectroscopy

The g-values of the Rieske cluster in different proteins reveal subtle variations. Generally, the Oxy signal is wider and characterized by a lower $g_x$ value than the Fd signal. The mutation β-L98K in CDO-Oxy did not alter the EPR signal of the Rieske cluster. The A50W mutation in NDO-Fd shows small changes in the g-values, suggesting conformational changes in the vicinity of the cluster.

Previous studies have reported EPR spectra of Rieske clusters from various Oxys and Fds[8–14]. The reported g-values are highly similar to those observed in this study (Table S7), with $g_z$ generally ranging from 2.01 to 2.03, $g_y$ from 1.90 to 1.91 and $g_x$ from 1.45 to 1.79. Previously reported EPR spectra of Rieske cluster containing Fds are more sparse, but they show the same trend of a less anisotropic EPR signal. The $g_y$ value is slightly lower, and the $g_x$ value is slightly higher than for the Oxys. For example, the previously reported $g_{zyx}$ values for TDO-Fd are 2.01, 1.86, and 1.81[15].

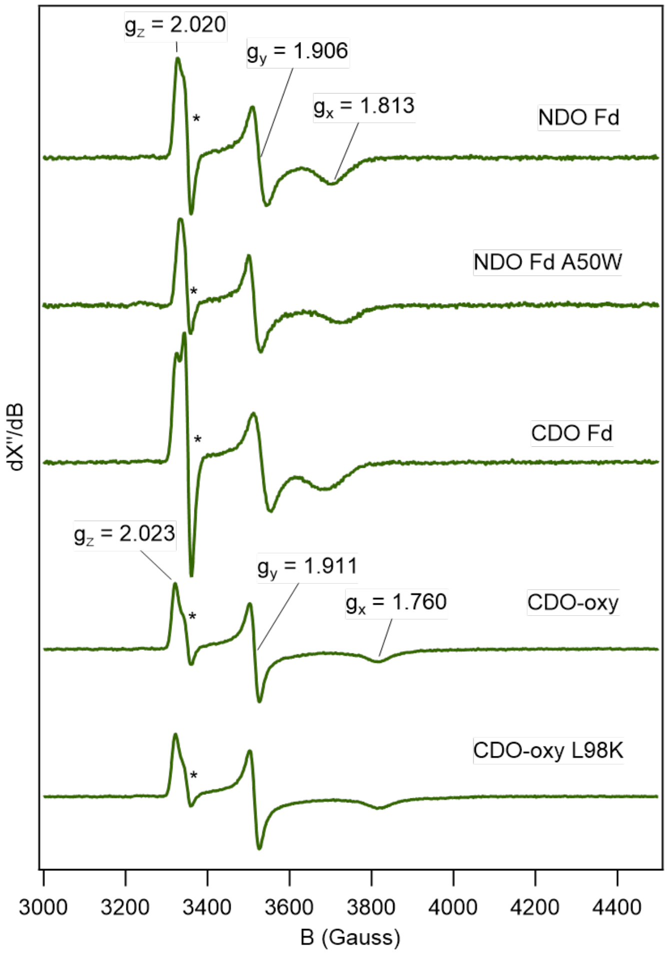

**Figure S21.** EPR of the Rieske [2Fe-2S] clusters of the different ferredoxin and oxygenase variants. The spectra were obtained from samples poised at −213 mV (NDO-Fd), −283 mV (NDO-Fd A50W), −202 mV (CDO-Fd), −212 mV (CDO-Oxy) and −228 mV (CDO-Oxy L98K). *Sharp isotropic signal indicative of an organic radical that is attributed to one of the redox dyes. All spectra have been normalized for the amplitude of the signal at $g_y$. EPR parameters are given in Table S7.

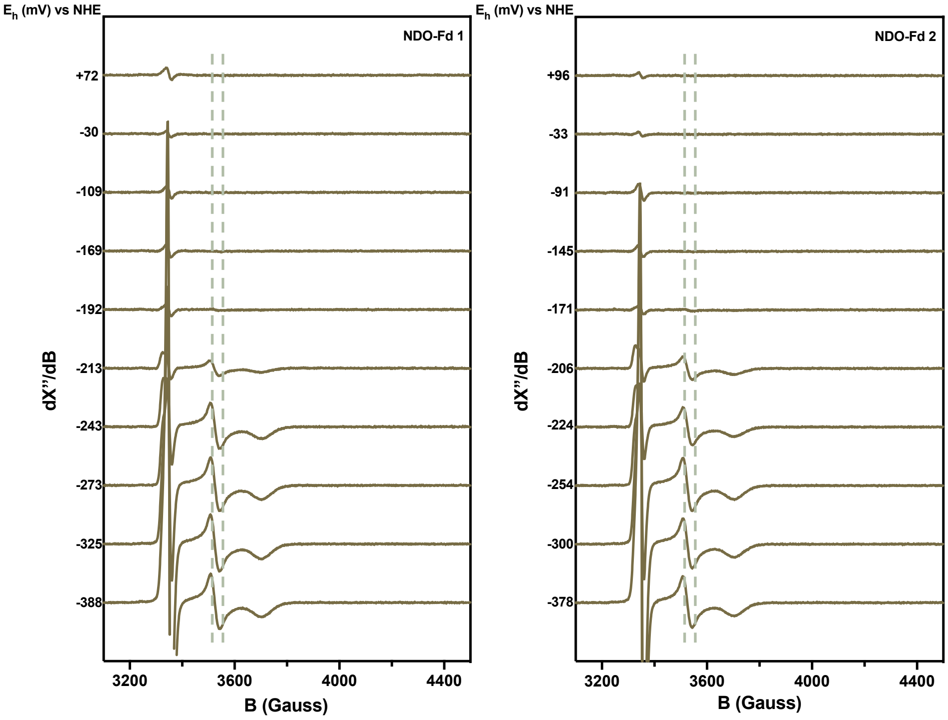

**Figure S22.** EPR spectroscopy of the Rieske cluster in NDO-Fd during titration experiments, performed in duplicate.

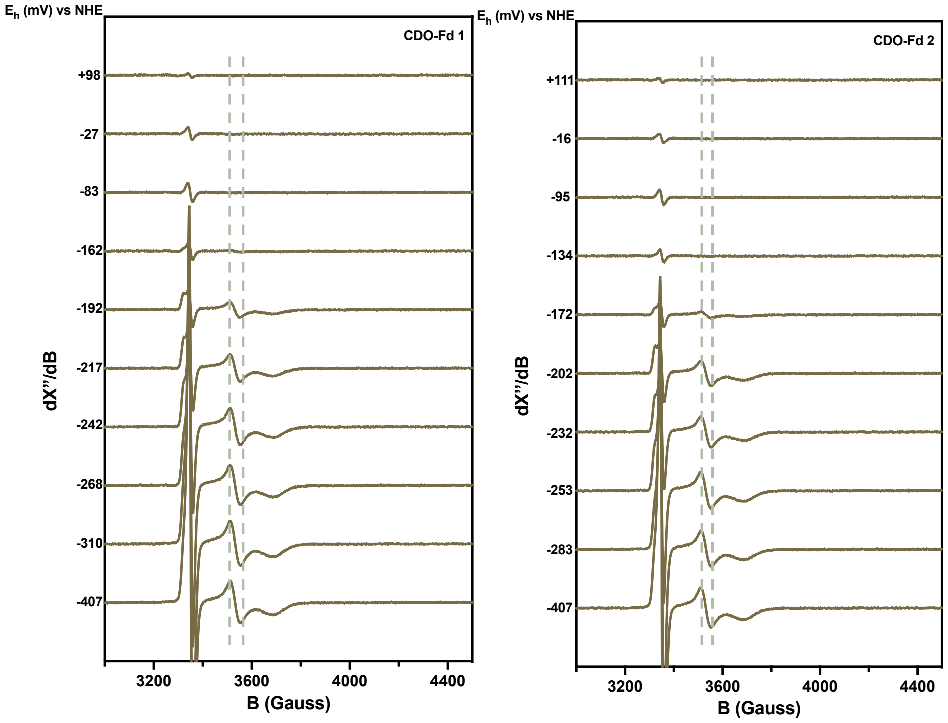

**Figure S23.** EPR spectroscopy of the Rieske cluster in CDO-Fd during titration experiments, performed in duplicate.

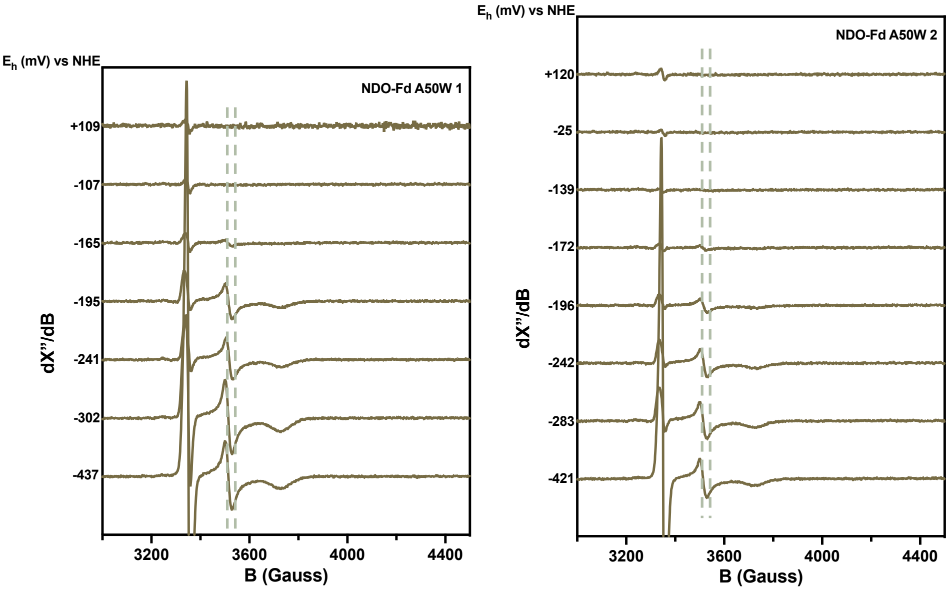

**Figure S24.** EPR spectroscopy of the Rieske cluster in NDO-Fd A50W during titration experiments, performed in duplicate.

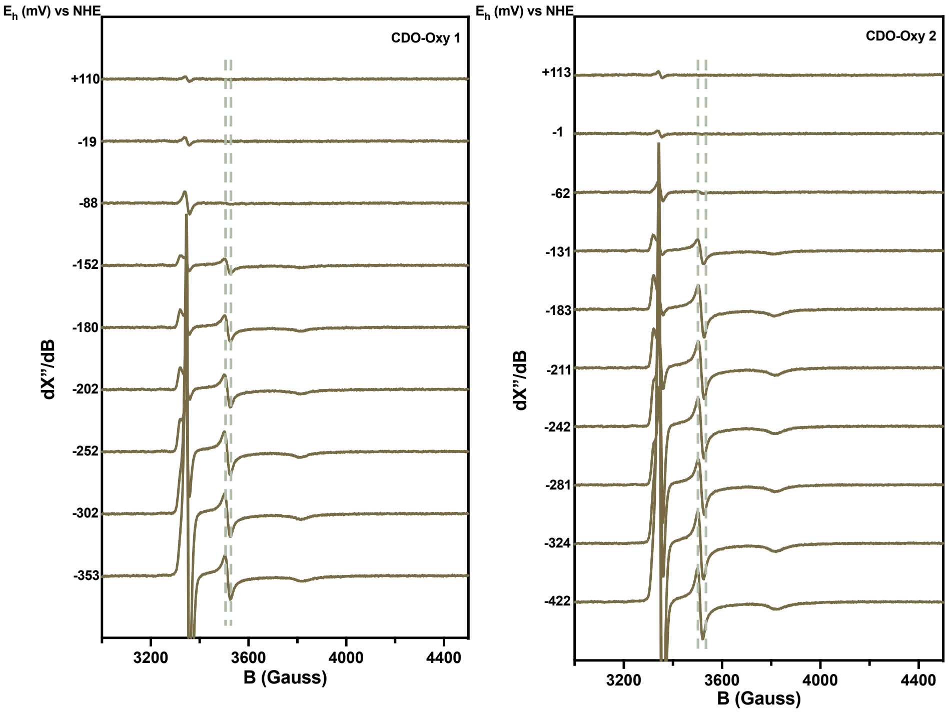

**Figure S25.** EPR spectroscopy of the Rieske cluster in CDO-Oxy during titration experiments, performed in duplicate.

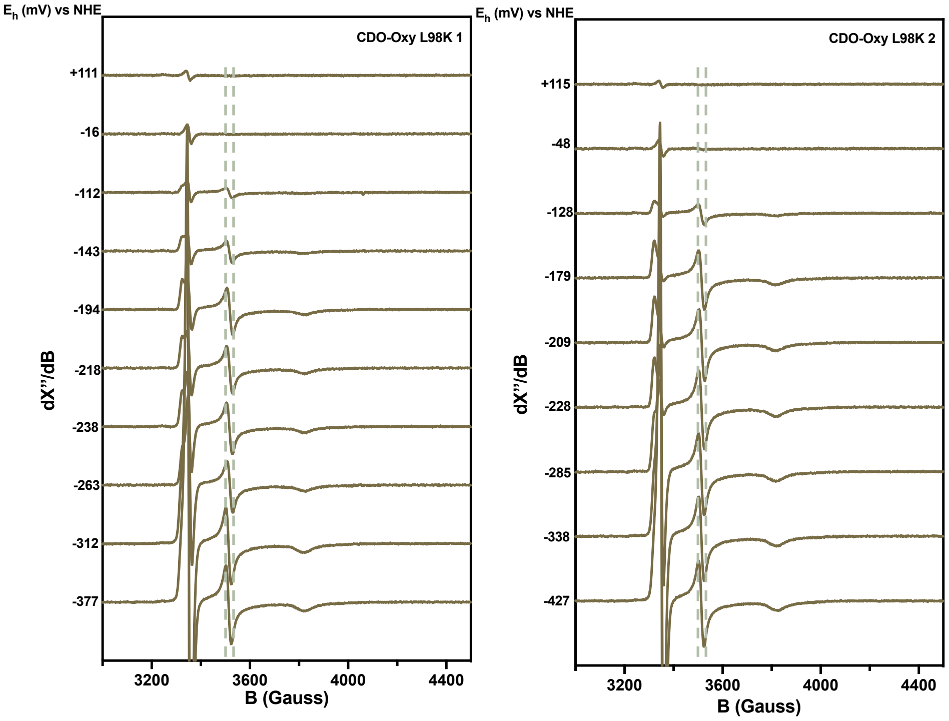

**Figure S26.** EPR spectroscopy of the Rieske cluster in CDO-Oxy L98K during titration experiments, performed in duplicate.

#### 6. MST reports

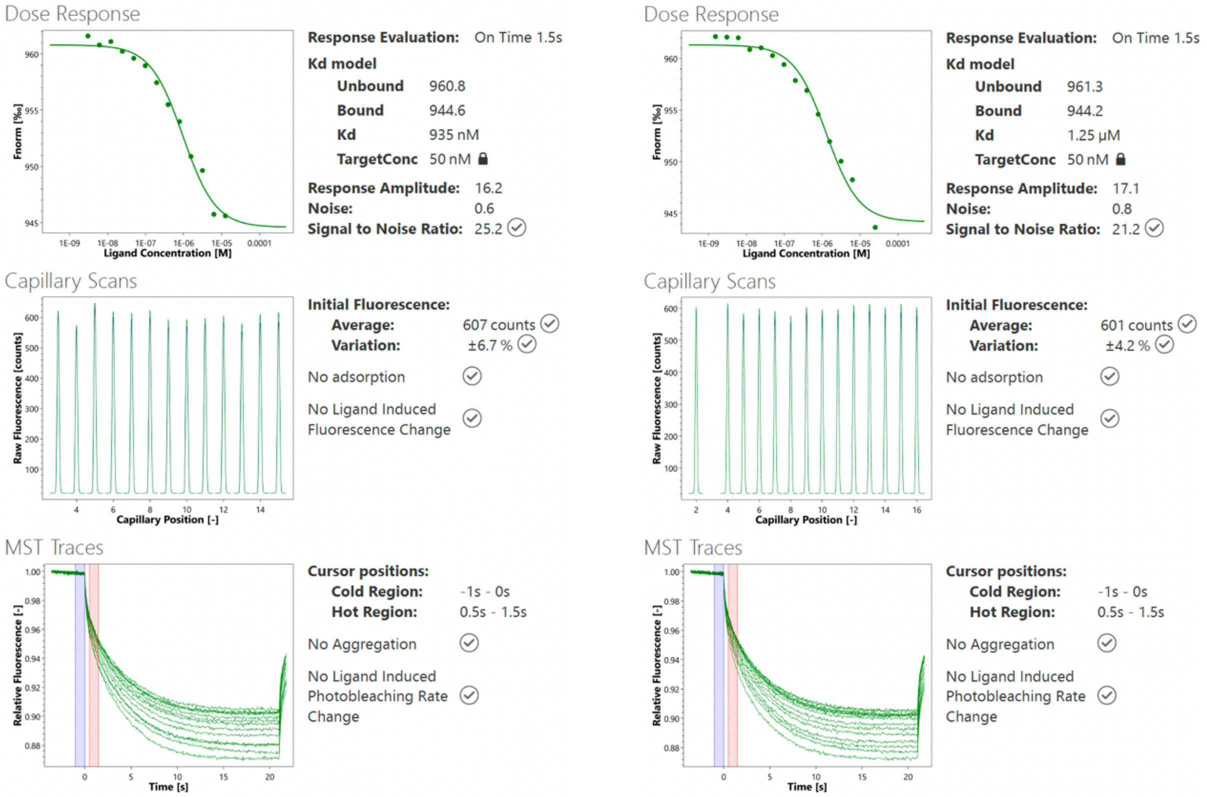

**Figure S27.** MST report of ligand NDO-Fd A50W binding to target CDO-Oxy WT, performed in duplicate.

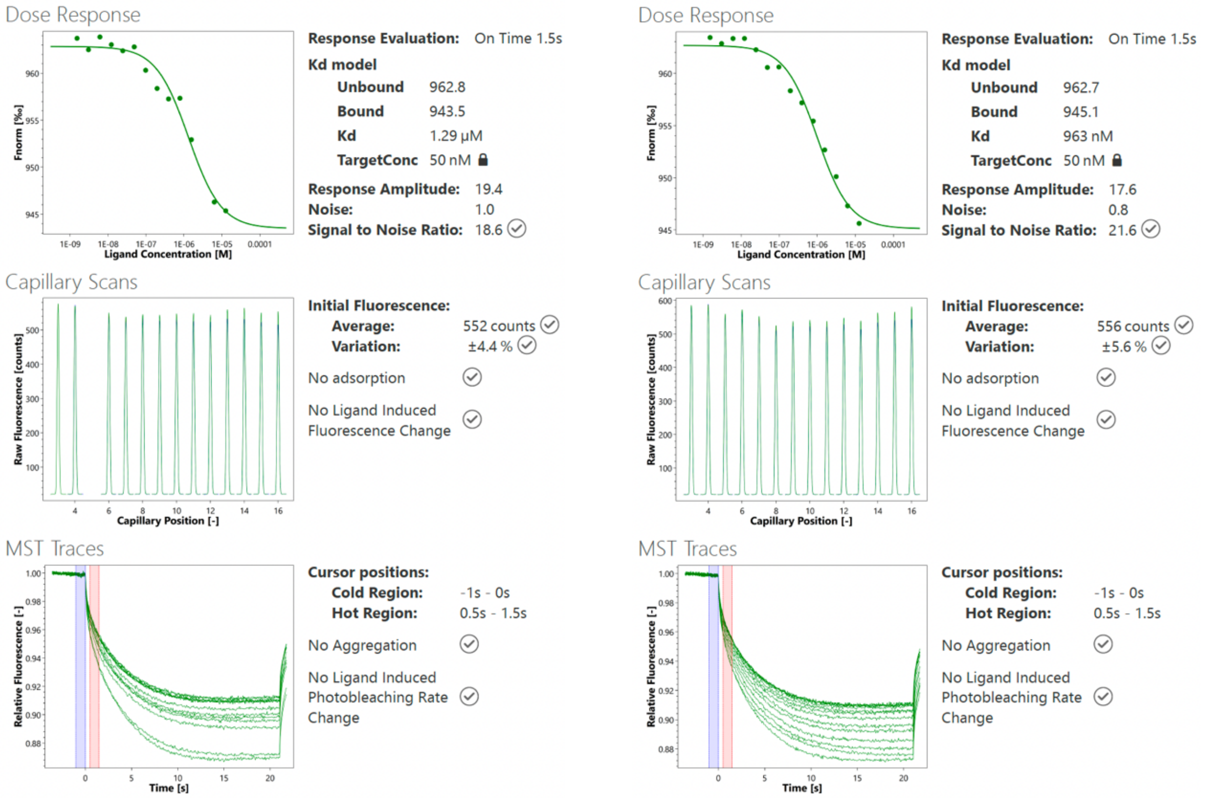

**Figure S28.** MST report of ligand NDO-Fd A50W binding to target CDO-Oxy L98K, performed in duplicate.

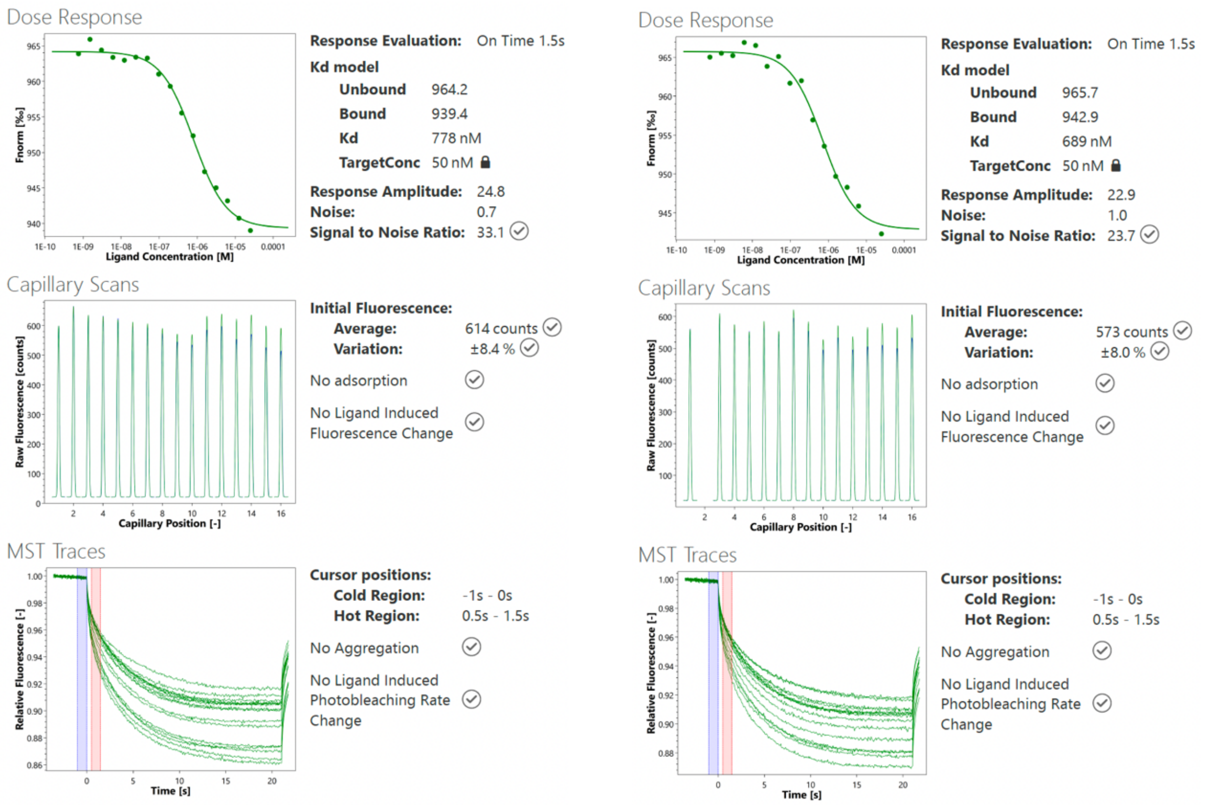

**Figure S29.** MST report of ligand CDO-Fd binding to target CDO-Oxy WT, performed in duplicate.

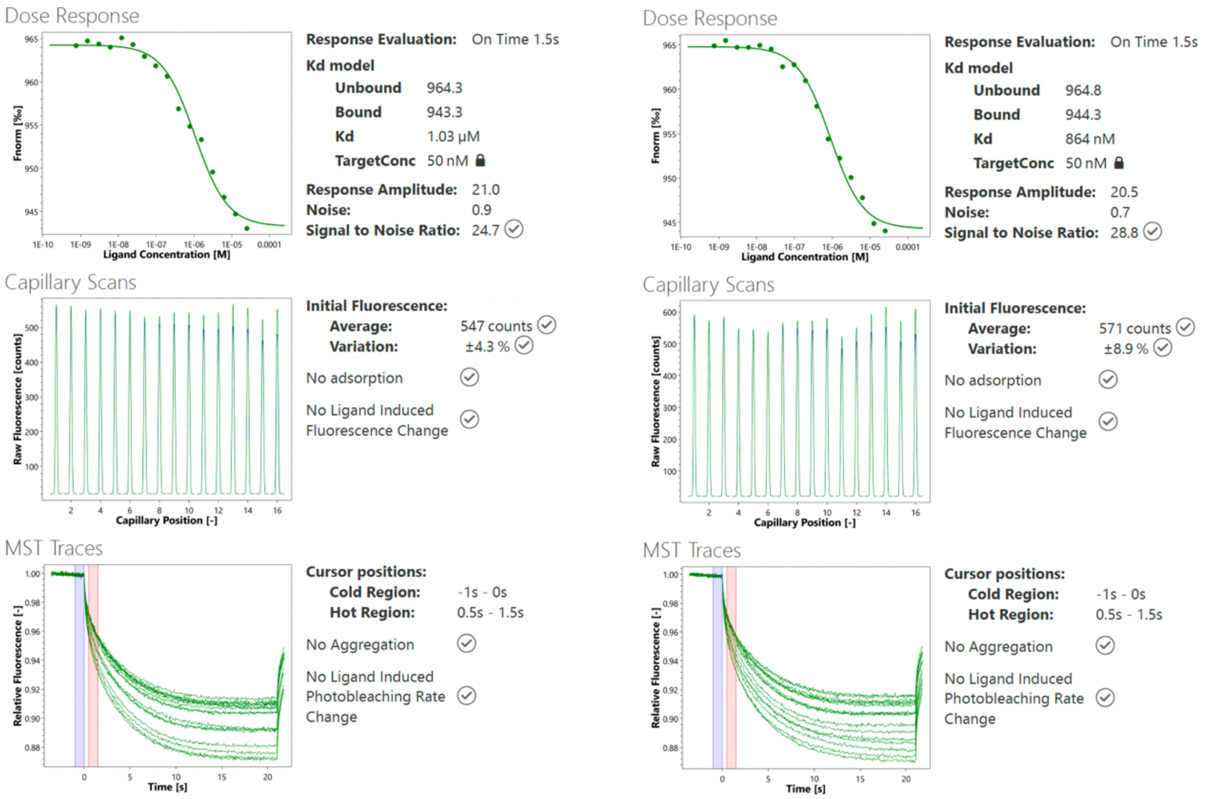

**Figure S30.** MST report of ligand NDO-Fd binding to target CDO-Oxy WT, performed in duplicate.

#### 7. Sequences of proteins used or generated in this work

>CDO-Red (UniProt ID: Q51747)
MIKSIVIIGAGLAGATATRYLRAQGYQGKIHLVGEELHVAYDRPSLSKDTLSGKVVE
PPAILDPCWYASADIDLHLGVRVTGIDVVNHQVLFESGDILAYDRLLLATGARARRM
AITGSELAGIHTLRDRADSQALRQALEPGQSLVIVGGGLIGCEVATTAINAGAHVTVL
EAGDELLLRVLGRSTGAWCRNELERLGVRVELNAQAAHFEGEGHVHAVVCADGRR
IAAGTVLVSIGAEPADELARAAGIACERGVVVDATGASSCPAVFAAGDVAAWPLRS
GELRSLETYLNSHMQAETAAAAMLGKSIPALQVPTSWTEIAGHRIQMVGDIEGPGEV
VLRGNVENGQPLVQFRVLDGRVEAATAINAPEDFPVATRLVADHIPVSATKLQDASS
NLRDFMKAKAERCE*

>CDO-Fd (UniProt ID: Q51746)
MTFSKVCEVSDVPVGDALQVESKGEAVAIFNVDGELFATQDRCTHGDWSLSEGGYL
EGDIVECSLHMGRFCVRTGKVKAAPPCEPLKIYPIRIDGSDVFVDFDAGYLAP*

>CDO-Oxy (UniProt ID: Q51743, Q51744)
MSSIINKEVQEAPLKWVKNWSDEEIKALVDEEKGLLDPRIFSDQDLYEIELERVFARS
WLLLGHEGHIPKAGDYLTTYMGEDPVIVVRQKDRSIKVFLNQCRHRGMRIERSDFG
NAKSFTCTYHGWAYDTAGNLVNVPYEKEAFCDKKEGDCGFDKADWGPLQARVDT
YKGLIFANWDTEAPDLKTYLSDATPYMDVMLDRTEAVTQVITGMQKTVIPCNWKF
AAEQFCSDMYHAGTMAHLSGVLSSLPPEMDLSQVKLPSSGNQFRAKWGGHGTGWF
NDDFALLQAIMGPKVVDYWTKGPAAERAKERLGKVLPADRMVAQHMTIFPTCSFLP
GINTVRTWHPRGPNEIEVWSFIVVDADAPEDIKEEYRRKNIFTFNQGGTYEQDDGEN
WVEVQRGLRGYKARSRPLCAQMGAGVPNKNNPEFPGKTSYVYSEEAARGFYHHWS
RMMSEPSWDTLKS*
MTSADLTKPIEWPEMPVSLELQNAVEQFYYREAQLLDYQNYEAWLALLTQDIQYW
MPIRTTHTSRNKAMEYVPPGGNAHFDETYESMRARIRARVSGLNWTEDPPSRSRHIV
SNVIVRETESAGTLEVSSAFLCYRNRLERMTDIYVGERRDILLRVSDGLGFKIAKRTIL
LDQSTITANNLSQFF*

>NDO-Red (UniProt ID: Q52126)
MELLIQPNNRIIPFSAGANLLEVLRENGVAISYSCLSGRCGTCRCRVIDGSVIDSGAEN
GQSNLTDKQYVLACQSVLTGNCAIEVPEADEIVTHPARIIKGTVVAVESPTHDIRRLR
VRLSKPFEFSPGQYATLQFSPEHARPYSMAGLPDDQEMEFHIRKVPGGRVTEYVFEH
VREGTSIKLSGPLGTAYLRQKHTGPMLCVGGGTGLAPVLSIVRGALKSGMTNPILLY
FGVRSQQDLYDAERLHKLAADHPQLTVHTVIATGPINEGQRAGLITDVIEKDILSLAG
WRAYLCGAPAMVEALCTVTKHLGISPEHIYADAFYPGGI*

>NDO-Fd (UniProt ID: P0A185)
MTVKWIEAVALSDILEGDVLGVTVEGKELALYEVEGEIYATDNLCTHGSARMSDGY
LEGREIECPLHQGRFDVCTGKALCAPVTQNIKTYPVKIENLRVMIDLS*

>NDO-Oxy (UniProt ID: P0A110, P0A112)
MNYNNKILVSESGLSQKHLIHGDEELFQHELKTIFARNWLFLTHDSLIPAPGDYVTAK
MGIDEVIVSRQNDGSIRAFLNVCRHRGKTLVSVEAGNAKGFVCSYHGWGFGSNGEL
QSVPFEKDLYGESLNKKCLGLKEVARVESFHGFIYGCFDQEAPPLMDYLGDAAWYL
EPMFKHSGGLELVGPPGKVVIKANWKAPAENFVGDAYHVGWTHASSLRSGESIFSS
LAGNAALPPEGAGLQMTSKYGSGMGVLWDGYSGVHSADLVPELMAFGGAKQERL
NKEIGDVRARIYRSHLNCTVFPNNSMLTCSGVFKVWNPIDANTTEVWTYAIVEKDM
PEDLKRRLADSVQRTFGPAGFWESDDNDNMETASQNGKKYQSRDSDLLSNLGFGED
VYGDAVYPGVVGKSAIGETSYRGFYRAYQAHVSSSNWAEFEHASSTWHTELTKTTD
R*
MMINIQEDKLVSAHDAEEILRFFNCHDSALQQEATTLLTQEAHLLDIQAYRAWLEHC
VGSEVQYQVISRELRAASERRYKLNEAMNVYNENFQQLKVRVEHQLDPQNWGNSP
KLRFTRFITNVQAAMDVNDKELLHIRSNVILHRARRGNQVDVFYAAREDKWKRGEG
GVRKLVQRFVDYPERILQTHNLMVFL*

> Fd-CN$_R$C
MTFSKVCEVSDVPVGDALQVESKGEAVAIFNVDGELFATQNLCTHGSARMSDGYLE
GREIECPLHQGRFDVCTGKVKAAPPCEPLKIYPIRIDGSDVFVDFDAGYLAP*

> Fd-NC$_R$N
MTVKWIEAVALSDILEGDVLGVTVEGKELALYEVEGEIYATDDRCTHGDWSLSEGG
YLEGDIVECSLHMGRFCVRTGKALCAPVTQNIKTYPVKIENLRVMIDLS*

>TDO-Red (UniProt ID: A5W4E9)
MATHVAIIGNGVGGFTTAQALRAEGFEGRISLIGDEPHLPYDRPSLSKAVLDGSLERP
PILAEADWYGEARIDMLTGPEVTALDVQTRTISLDDGTTLSADAIVIATGSRARTMAL
PGSQLPGVVTLRTYGDVQVLRDSWTSATRLLIVGGGLIGCEVATTARKLGLSVTILE
AGDELLVRVLGRRIGAWLRGLLTELGVQVELGTGVVGFSGEGQLEQVMASDGRSFV
ADSALICVGAEPADQLARQAGLACDRGVIVDHCGATLAKGVFAVGDVASWPLRAG
GRRSLETYMNAQRQAAAVAAAILGKNVSAPQLPVSWTEIAGHRMQMAGDIEGPGD
FVSRGMPGSGAALLFRLQERRIQAVVAVDAPRDFALATRLVEARAAIEPARLADLSN
SMRDFVRANEGDLT*

>TDO-Fd (UniProt ID: A5W4F0)
MTWTYILRQGDLPPGEMQRYEGGPEPVMVCNVDGEFFAVQDTCTHGDWALSDGYL
DGDIVECTLHFGKFCVRTGKVKALPACKPIKVFPIKVEGDEVHVDLDNGELK*

>TDO-Oxy (UniProt ID: A5W4F2, A5W4F1)
MNQTDTSPIRLRRSWNTSEIEALFDEHAGRIDPRIYTDEDLYQLELERVFARSWLLLG
HETQIRKPGDYITTYMGEDPVVVVRQKDASIAVFLNQCRHRGMRICRADAGNAKAF
TCSYHGWAYDTAGNLVNVPYEAESFACLNKKEWSPLKARVETYKGLIFANWDENA
VDLDTYLGEAKFYMDHMLDRTEAGTEAIPGVQKWVIPCNWKFAAEQFCSDMYHA
GTTSHLSGILAGLPEDLEMADLAPPTVGKQYRASWGGHGSGFYVGDPNLMLAIMGP
KVTSYWTEGPASEKAAERLGSVERGSKLMVEHMTVFPTCSFLPGINTVRTWHPRGP

NEVEVWAFTVVDADAPDDIKEEFRRQTLRTFSAGGVFEQDDGENWVEIQHILRGHK
ARSRPFNAEMSMDQTVDNDPVYPGRISNNVYSEEAARGLYAHWLRMMTSPDWDA
LKATR*
MIDSANRADVFLRKPAPVAPELQHEVEQFYYWEAKLLNDRRFEEWFALLAEDIHYF
MPIRTTRIMRDSRLEYSGSREYAHFDDDATMMKGRLRKITSDVSWSENPASRTRHLV
SNVMIVGAEAEGEYEISSAFIVYRNRLERQLDIFAGERRDTLRRNTSEAGFEIVNRTILI
DQSTILANNLSFFF*

#### References

1. Feyza Özgen, F. *et al.* Artificial light-harvesting complexes enable Rieske oxygenase catalyzed hydroxylations in non-photosynthetic cells. *Angew. Chem., Int. Ed.* **59**, 3982–3987 (2020).
2. Runda, M. E., Kremser, B., Özgen, F. F. & Schmidt, S. An optimized system for the study of Rieske oxygenase-catalyzed hydroxylation reactions in vitro. *ChemCatChem* **15**, e202300371 (2023).
3. Prats Luján, A., Bhat, M. F., Saravanan, T. & Poelarends, G. J. Chemo- and enantioselective photoenzymatic ketone reductions using a promiscuous flavin-dependent nitroreductase. *ChemCatChem* **14**, e202200043 (2022).
4. Onaran, M. B. & Seto, C. T. Using a lipase as a high-throughput screening method for measuring the enantiomeric excess of allylic acetates. *J. Org. Chem.* **68**, 8136–8141 (2003).
5. Jorgensen, A. D., Picel Kurt C. & Stamoudis Vassilis C. Prediction of gas chromatography flame ionization detector response factors from molecular structures. *Anal. Chem.* **62**, 683–689 (1990).
6. Madeira, F. *et al.* The EMBL-EBI job dispatcher sequence analysis tools framework in 2024. *Nucleic Acids Res.* **52**, W521–W525 (2024).
7. Bagnéris, C., Cammack, R. & Mason, J. R. Subtle difference between benzene and toluene dioxygenases of *Pseudomonas putida*. *Appl. Environ. Microbiol.* **71**, 1570–1580 (2005).
8. Quareshy, M. *et al.* Structural basis of carnitine monooxygenase CntA substrate specificity, inhibition, and intersubunit electron transfer. *J. Biol. Chem.* **296**, 100038 (2021).
9. Rogers, M. S. & Lipscomb, J. D. Salicylate 5-Hydroxylase: Intermediates in Aromatic Hydroxylation by a Rieske Monooxygenase. *Biochemistry* **58**, 5305–5319 (2019).
10. Lee, J., Simurdiak, M. & Zhao, H. Reconstitution and characterization of aminopyrrolnitrin oxygenase, a Rieske *N*-oxygenase that catalyzes unusual arylamine oxidation. *J. Biol. Chem.* **280**, 36719–36728 (2005).
11. Coulter, E. D., Moon, N., Batie, C. J., Dunham, W. R. & Ballou, D. P. Electron paramagnetic resonance measurements of the ferrous mononuclear site of phthalate dioxygenase substituted with alternate divalent metal ions: Direct evidence for ligation of two histidines in the copper(II)- reconstituted protein. *Biochemistry* **38**, 11062–11072 (1999).
12. Wolfe, M. D., Parales, J. V., Gibson, D. T. & Lipscomb, J. D. Single turnover chemistry and regulation of O2 activation by the oxygenase component of naphthalene 1,2-dioxygenase. *J. Biol. Chem.* **276**, 1945–1953 (2001).
13. Shanmugam, M., Quareshy, M., Cameron, A. D., Bugg, T. D. H. & Chen, Y. Light-Activated Electron Transfer and Catalytic Mechanism of Carnitine Oxidation by Rieske-Type Oxygenase from Human Microbiota. *Angew. Chem., Int. Ed.* **60**, 4529–4534 (2021).
14. Baratto, M. C. *et al.* Spectroscopic characterisation of the naphthalene dioxygenase from *Rhodococcus* sp. Strain NCIMB12038. *Int. J. Mol. Sci.* **20**, (2019).
15. Subramanian, V., Liu, T. N. & Yeh, W. K. Purification and properties of ferredoxin(TOL): A component of toluene dioxygenase from *Pseudomonas putida* F1. *J. Biol. Chem.* **260**, 2355–2363 (1985).
